# Beta turn propensity and a model polymer scaling exponent identify disordered proteins that phase separate

**DOI:** 10.1101/2020.07.06.189613

**Authors:** Elisia A. Paiz, Jeffre H. Allen, John J. Correia, Nicholas C. Fitzkee, Loren E. Hough, Steven T. Whitten

## Abstract

The complex cellular milieu can spontaneously de-mix in a process controlled in part by proteins that are intrinsically disordered (ID). A protein’s propensity to de-mix is thought to be driven by the preference for protein-protein rather than protein-solvent interactions. The hydrodynamic size of monomeric proteins, as quantified by the polymer scaling exponent (*v*), is driven by a similar balance. We hypothesize that mean *v*, as predicted by the protein sequence, will be smaller for proteins with a strong propensity to de-mix. To test this hypothesis, we analyzed protein databases containing subsets that are either folded, disordered, or disordered and known to spontaneously phase separate. We find that the phase separating disordered proteins, on average, have lower calculated values of *v* compared to their non-phase separating counterparts. Moreover, these proteins have a higher sequence-predicted propensity for β-turns. Using a simple, surface area-based model, we propose a physical mechanism for this difference: transient β-turn structures reduce the desolvation penalty of forming a protein-rich phase and increase exposure of atoms involved in π/sp^2^ electronic interactions. By this mechanism, β-turns act as energetically favored nucleation points, which may explain the increased propensity for turns in ID regions (IDRs) that are utilized biologically for phase separation. Phase separating IDRs, non-phase separating IDRs, and folded regions could be distinguished by combining *v* and β-turn propensity, and we propose a new algorithm, ParSe (<u>par</u>tition <u>se</u>quence), for predicting phase separating protein regions. ParSe is able to accurately identify folded, disordered, and phase-separating protein regions from the primary sequence.

## Introduction

Protein liquid-liquid phase separation (LLPS) is increasingly recognized as an important organizing phenomenon in cells. LLPS is a reversible process whereby complex protein mixtures spontaneously de-mix into liquid droplets that are enriched in a particular protein; concomitantly, surrounding regions are depleted of that protein (1). This de-mixing transition is thought to provide temporal and spatial control over intracellular interactions by assembling collections of proteins into structures called membraneless organelles (2), a key step in the regulatory function of P bodies, the nucleolus, and germ granules (3–5). The physical mechanisms responsible for LLPS are not fully understood, but it is known to be facilitated primarily, though not exclusively (6, 7), by proteins that are intrinsically disordered (ID) or contain large ID regions (IDRs) (8, 9, 2) – proteins termed intrinsically disordered proteins (IDPs). The propensity for a particular protein to phase separate is, in general, determined by the balance of intra-molecular and solvent interactions. In part, based on mechanistic insights into the nature of these interactions, several groups have developed sequence-based predictors to identify LLPS regions (10–12).

The same interactions that drive LLPS have also been hypothesized to affect hydrodynamic size of monomeric IDRs. The hydrodynamic size of IDRs has been found to vary substantially with the primary sequence (13, 14) and appears important for the function of IDRs. For example, some IDPs regulate the remodeling of cellular membranes, and their size controls curvature at membrane surfaces (15, 16). Conceptually, favorable interactions with the solvent give rise to mean ensemble dimensions for the polymer that are elongated and swollen when compared to the compacted dimensions observed when self-interactions dominate. One framework to quantify this relationship is derived from polymer theories developed for long homopolymers (17, 18). Despite some limitations (19), homopolymer theories have been successful in describing the properties of short, heteropolymeric IDRs (20–25). In particular, the polymer scaling exponent, *v*, has been used to extract information on the balance of protein-self and protein-solvent interactions (26, 27, 13). This exponent is obtained experimentally from the dependence of size (e.g., hydrodynamic radius, *R_h_*, or radius of gyration, *R_g_*) on polymer length, *N*, in the power law relationship, *R_h_* ∝ *N^v^*.

Because biomolecular LLPS includes the exchange of macromolecule-solvent interactions for macromolecule-macromolecule interactions (28–30), *v* could be a predictor of LLPS potential among heteropolymeric IDRs (19, 24, 25, 31). Numerous studies have already found that the hydrodynamic dimensions of some IDRs are correlated to the temperature dependence of the demixing transition (24, 32, 33). Moreover, Dignon and coworkers found in molecular simulations that the critical temperature of phase separation and the internal scaling exponent, *v_int_*, which is a variation on *v* calculated as the average intrachain pairwise distance in a single chain, 〈*R_i-j_*〉 ∝ |*i* – *j*|*^ν_int_^* (34), are correlated properties (19). However, whether the scaling exponent of a monomeric IDR can predict its potential for LLPS remains unclear.

We (14, 22, 35) and others (13, 36) have developed sequence-based methods to predict the hydrodynamic dimensions of IDPs, allowing us to test whether an IDP’s potential to phase separate can be predicted from its monomeric scaling exponent, *v*. We hypothesize that the same self-interactions that facilitate LLPS will reduce the mean *R_h_* (and thus *v*) for IDRs competent to phase separate into protein-rich droplets when compared to IDRs that are not (19, 24, 25, 31). Indeed, this trend is evident and shown schematically in Figure 1. Moreover, we find that β-turn propensity (37–39) is higher for phase separating IDRs, and we develop a simple surface area-based approach to show how β-turns can reduce the desolvation penalty in LLPS. Using these observations, we developed the computer algorithm ParSe (<u>par</u>tition <u>se</u>quence) for predicting folded, phase-separating ID, and non-phase-separating ID regions given only the primary sequence. A basic version of the algorithm has been made web-accessible at: folding.chemistry.msstate.edu/utils/ parse.html. We find that the predictions from ParSe had strong correlations to other predictors of protein phase separation (10–12, 40), indicating that β-turns and *v* may provide physically meaningful insight into the diverse molecular mechanisms driving LLPS.

**Figure 1.**
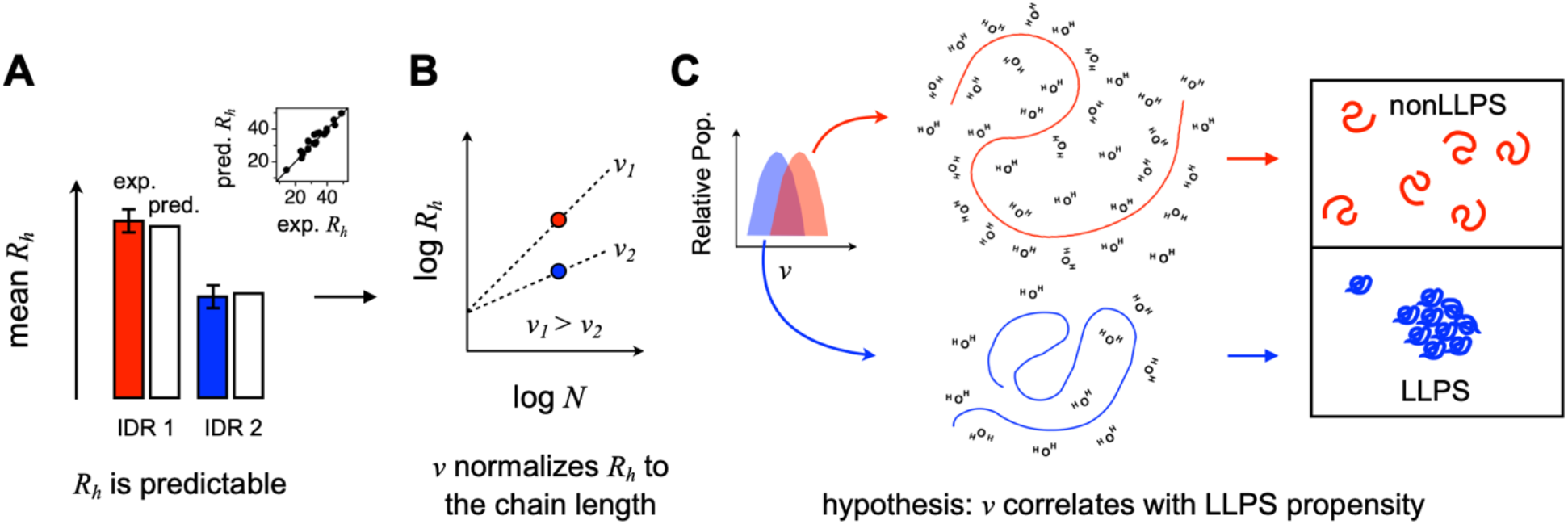
IDR propensity for LLPS predicted from hydrodynamic size. **A)** The mean *R_h_* determined experimentally for monomeric IDPs (or excised IDRs) can be predicted from sequence. The inset compares predicted to observed for a set of 23 IDPs (listed in Table S1). *R_h_* is in Å. A full-scale version of this inset is in Supporting Information, Fig. S1. **B)** Converting sequence calculated *R_h_* to *v* normalizes the hydrodynamic size to the protein chain length. **C)** IDRs that strongly prefer interactions with the solvent (cartoon shows waters) over self are likely to exhibit swollen structures and remain monomeric rather than drive transitions to protein-rich phase separated states. The inset in this panel is based on *v* distributions calculated in two different sequence sets; a full-scale version of the inset is in Supporting Information, Fig. S2.

## Results

### Sequence calculated polymer scaling exponent, *v_model_*, is reduced in IDRs from proteins that exhibit LLPS when compared to IDRs in general

Proteins have modular structures, which can consist of folded regions and IDRs. Among the IDRs, some potentially drive phase separation, and some do not. For example, the 685-residue Sup35 protein from yeast has three domains (41); the ID N-terminal prion domain (residues 1-124), the ID middle domain (residues 125-254), and the folded (42) C-terminal catalytic domain (residues 255-685). Of these domains, only the N-terminal prion domain mediates phase separation (41). Here, the word *domain* is used to identify a protein *region* that has distinctive features or properties, and not necessarily to indicate a globular structure (41, 43). For the present work, we use *domain* and *region* interchangeably in this manner, though with a preference for using *region*. The goal of this work is to determine if IDRs that drive phase separation show differences in predicted hydrodynamic dimensions of their equilibrium conformational ensembles as compared to IDRs that do not (19, 24, 25, 31). Specifically, we hypothesize that when excised from the parent protein, compacted ensemble dimensions would indicate high LLPS potential for the disordered polypeptide, while elongated sizes would indicate low LLPS potential (Figure 1).

To test this idea against large numbers of proteins, we used a sequence-based calculation of the radius of hydration (*R_h_*) that has been found to be accurate for monomeric IDPs (14, 35, 44). Figure S1A shows predicted and experimental mean *R_h_* for a set of 23 IDPs (35, 44–61), demonstrating overall good agreement. This IDP set is identified in Table S1. The sequence calculated *R_h_* uses the net charge and intrinsic chain propensity for the polyproline II backbone conformation (see Experimental Procedures), both known to promote elongated hydrodynamic dimensions in disordered ensembles (13, 14). To normalize *R_h_* to the chain length, and distinguish compacted versus elongated predicted dimensions, we converted the sequence calculated value to a polymer scaling exponent by,

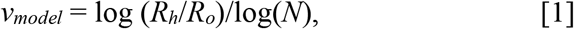

where *N* is the number of residues, and *R_O_* is 2.16 Å obtained from simulated IDP ensembles (62). Experimental *R_O_* for the set of 23 IDPs is ~2.1 Å (Fig. S1B), showing good agreement. For these 23 IDPs, which are not known to phase separate and thus referred to as the null set, the mean ± σ (standard deviation) in *v_model_* was 0.558 ± 0.019 (Table 1).

**Table 1.** Summary of mean *v_model_* in the protein sequence sets.

| | # sequences | $v_{model}^a$ | z-test w/ null <sup>b</sup> | z-test w/ folded <sup>b</sup> |
| --- | --- | --- | --- | --- |
| null set | 23 | $0.558 \pm 0.019$ | 0.5 | 8.8e-08 |
| testing set | 224 | $0.542 \pm 0.020$ | 8.2e-05 | 2.7e-04 |
| folded set | 82 | $0.536 \pm 0.008$ | 8.8e-08 | 0.5 |

<sup>a</sup> mean ± standard deviation
<sup>b</sup> one-tail p-value calculated using the set variances

Next, we compiled a list of IDRs from proteins annotated to phase separate. We began with the IDRs from 43 proteins verified to exhibit phase separation behavior *in vitro*, previously collated by Vernon *et al* (10). To this set, we added IDRs from 59 human proteins listed in the PhaSePro database as showing LLPS behavior (63), and 18 IDRs annotated “liquid-liquid phase separation” (IDPO:00041) in the DisProt database (64). Duplicate entries from merging the three sources were removed, as were IDRs with *N*<20. To identify IDRs in the Vernon *et al* protein set, we used the GeneSilico MetaDisorder service that predicts the presence of ID in a sequence (65). The PhaSePro database already annotates each entry with its predicted IDRs from using IUPred2 (66), which we kept for this analysis. Because DisProt is manually curated for verified cases of ID, we assumed LLPS annotated IDRs in DisProt lacked folded regions (i.e., each was fully ID). In total, this resulted in 224 IDRs from proteins with known phase separation behavior. The IDRs are identified in Table S2. This set was designated as the testing set. Trends identified in the testing set were used to analyze the entire human proteome, the full DisProt database, and the full PhaSePro database (see below).

On average, *v_model_* was reduced in the IDRs from known LLPS proteins, compared to the null set (i.e., the 23 IDPs not known to phase separate). The mean *v_model_* was 0.542 ± 0.020 for the testing set IDRs (Table 1). The *v_model_* distribution overlapped between the two sets (Fig. S2C), testing and null. Possibly contributing to the statistical overlap in *v_model_* between the two sets, most IDRs in the testing set have not been verified to drive phase separation, suggesting the set may contain some that do not. Like Sup35 and its two IDRs, only one of which directly mediates a de-mixing transition (41), proteins in the testing set could have IDRs not necessary for LLPS. Indeed, most testing set proteins have >1 predicted IDR (Table S2). Or, simply, the overlap could be a consequence of the small difference in means. Despite this unknown, a two-sample z-test using the variances gave a one-tail p-value of 8.2e-05 (Table 1), indicating the two sets are statistically different in mean *v_model_*. Though the distribution of *v_model_* values in both sets were similar to normal (Figs. S2A and S2B), the non-parametric Mann-Whitney U test that does not assume normal distributions was also used and likewise shows the null and testing sets as statistically different in mean *v_model_* (Table S3).

We sought to determine if we could distinguish IDRs known to drive LLPS from folded regions. In order to compile a list of folded regions, we took folded regions from the same proteins as we had previously taken the IDR regions to form the testing set. We used the Protein Data Bank (67) to identify those regions (*N*≥20) in the testing set proteins that are verified to adopt stable, globular structures, finding 82 such regions (Table S2). Compared to the ID-based testing and null sets, the mean *v_model_* was slightly lower in this set of folded sequences and found to be 0.536 ± 0.008 (Table 1). It is important to note that the equation used to predict *R_h_* from sequence, and thus calculate *v_model_*, was developed for IDPs and not designed for use with folded proteins. For a folded protein, experimental *v* is often ~0.3 (26) rather than the calculated values for *v_model_* here that are >0.5. Sequence calculated *v_model_* determined by eq. [1] represents the mean hydrodynamic dimensions when the polypeptide is disordered and omits effects that are associated with hydrophobic collapse and folding. In concept, *v_model_* approximates the dimensions of the unfolded chain prior to collapse or the formation of cooperative units of structure.

### Sequence calculated β-turn propensity is elevated in IDRs from proteins that exhibit LLPS

Because *v_model_* was calculated from the primary sequence, we determined the differences in composition between the testing, null, and folded sets. Particularly, we were interested in identifying those amino acid types that are either depleted or enriched in the testing set when compared to the null and folded sets. For example, the branched amino acids, I, L, and V, are each depleted in the testing set, whereas P, G, and S are enriched, relative to both the other sets (Fig. 2A). While additional amino acid types show depletion in the testing set compared to the folded and null sets (e.g., C and E) or enrichment, (e.g., N and Q), specific mention of I, L, V, P, G, and S is made for two reasons. First, I, L, and V have infrequent occurrence in the β-turn in surveys of stable, protein structures (37), while P, G, and S are preferred in this secondary structure type (38). As such, this result predicts the intrinsic propensity for β-turn is higher, on average, in the testing set when compared to the null and folded sets. Second, studies involving elastin-like polypeptide (ELP), a protein sequence known to undergo LLPS (68–70), have implicated transient β-turn structures in the protein-protein interactions that mediate condensate formation (71–73). The results shown in Figure 2A predict a role for the β-turn in protein-based LLPS that could be widespread in use and not limited to ELP-based systems.

**Figure 2.**
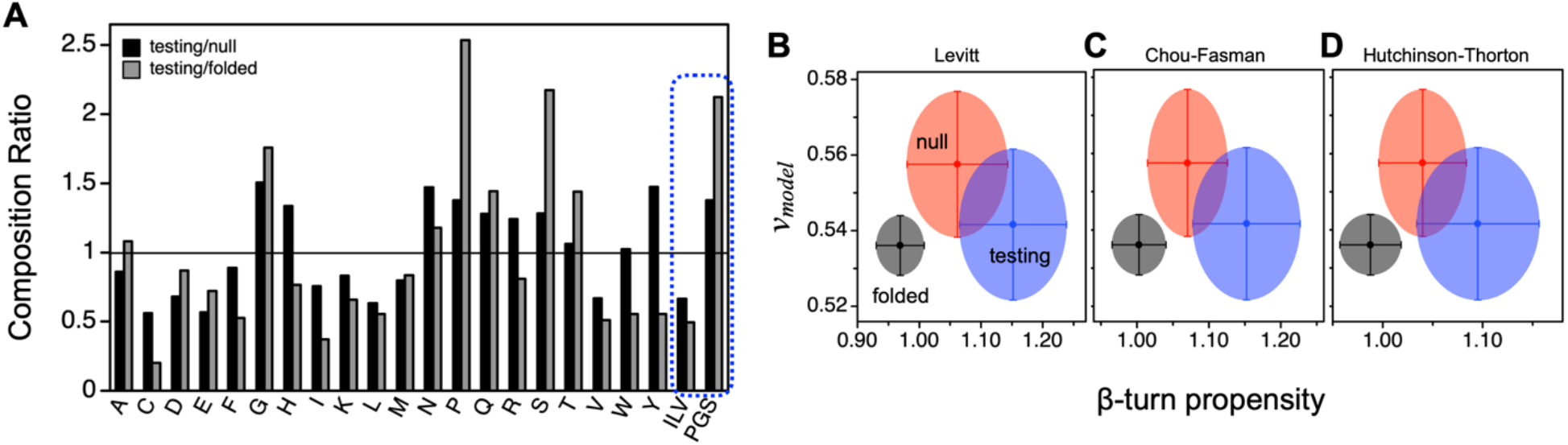
Composition differences among the sequence sets reveals that protein classes can be separated using *v_model_* and β-turn propensity. **A)** Composition ratio is the percent composition of each amino acid type, identified by its 1-letter code, in the training set divided by the percent composition in the null (black columns) and folded sets (gray columns). The two columns labeled ILV and PGS represent combining I+L+V and P+G+S content, respectively. Comparing sequence calculated *v_model_* in each set, testing (blue), null (red), and folded (black), to sequence calculated β-turn propensity using single position scales from **B)** Levitt (37) and **C)** Chou and Fasman (38), and **D)** four position scales from Hutchinson and Thorton (39). Data points and error bars show the mean and standard deviations in each set.

To determine if β-turn propensities are elevated in IDRs from LLPS proteins, we calculated the mean intrinsic β-turn propensity from sequence in the null, testing, and folded sets (Table 2). By using the amino acid scale of turn propensity developed by Levitt (37), which is reproduced in Table S4, we find the mean for the testing set was 1.152 ± 0.087. In comparison, the mean intrinsic turn propensity was lower in both the null and folded sets. A two-sample z-test using the variances indicated the mean values for intrinsic β-turn propensity were statistically different when compared between the three sets (Table 2); identical conclusions were obtained from the Mann-Whitney U test (Table S3).

**Table 2.** Summary of mean β-turn propensity in the protein sequence sets.

| | # sequences | β-turn propensity <sup>a</sup> | z-test w/ null <sup>b</sup> | z-test w/ folded <sup>b</sup> |
| --- | --- | --- | --- | --- |
| null set | 23 | $1.062 \pm 0.082$ | 0.5 | 6.4e-08 |
| testing set | 224 | $1.152 \pm 0.087$ | 2.6e-07 | 8.9e-142 |
| folded set | 82 | $0.969 \pm 0.039$ | 6.4e-08 | 0.5 |

<sup>a</sup> mean ± standard deviation, calculated using the scale from Levitt (37)
<sup>b</sup> one-tail p-value calculated using the set variances

### The three sets of protein regions, folded, null (IDRs) and testing (LLPS IDRs), form separate protein classes according to their mean values of *v_model_* and β-turn propensity

Figure 2B compares the mean *v_model_* of each set (i.e., testing, null, and folded) to the mean intrinsic β-turn propensity, showing that they typically occupy different regions in this plot. These results are robust to choice of scale that was employed for the sequence-based calculation of β-turn; the average intrinsic β-turn propensity was lowest in the folded set. For example, Figures 2C and 2D show identical results are obtained, whereby the mean intrinsic β-turn propensity is highest in the testing set, when the amino acid scale from Chou and Fasman is used instead (38), or when specificity for amino acid type in the four different turn positions (*i*, *i*+1, *i*+2, *i*+3) is used (39). The increased propensity for β-turn in the testing set is a curious result considering reverse turns are prevalent in globular structures (74) and often found at the protein surface (75) because it allows the polypeptide chain to fold back onto itself.

### β-turn structures reduce chain associated solvent waters, potentially driving intra-molecular contacts

To understand why β-turn structures might be associated with sequences that undergo LLPS, we investigated the balance of self and solvent interactions for this conformation. In lieu of large-scale molecular dynamics simulations, we considered the distribution of surface water molecules as captured by the conditional hydrophobic accessible surface area (CHASA). In the CHASA calculations, sterically-allowed solvent waters are placed near protein hydrophilic groups under the assumption that these waters will form hydrogen bonds with the peptide; then, hydrophobic surface area calculations are performed with these solvent waters present (76). We hypothesized that turn structures would facilitate protein-protein contacts because they had larger accessible hydrophobic patches even in the presence of bound solvent water molecules.

We randomly generated 1,000 structures containing β-turns and 1,000 non-turn structures of the ELP-derived peptide sequence GVPGVG (68–70), a sequence where transient β-turns have been implicated in self-association (71–73). Total accessible surface area (ASA), hydrophobic ASA, and CHASA are all lower in β-turn versus non-turn ensembles (Table S5), consistent with prior studies (72). Fewer hydration waters are associated with turn structures (37.1 ± 0.1 vs 44.4 ± 0.1 waters), meaning that the penalty for desolvation in LLPS will likely be lower for IDPs that preferentially sample the β-turn conformation. In addition, fixed conformations of β-turns also expose large contiguous regions of hydrophobic surface area relative to the random conformations (**Fig. 3**). The CHASA-placed solvation waters in GVPGVG are clustered when the peptide is in a β-turn conformation (**Fig. 3A, top**), exposing contiguous segments of hydrophobic accessible surface area for residues V2 and P3. Representative structures of non-turn conformations (**Fig. 3A, bottom**) do not exhibit these clusters. The contiguous hydrophobic segments present in β-turns may facilitate protein-protein association; two fixed β-turns can associate and bury a large relative fraction of hydrophobic accessible surface area (~110 Å^2^ per turn; based on docking with the GOLD software package). Finally, the π electrons sp^2^ hybridized atoms of the peptide bond (π/sp^2^ interactions) are thought to facilitate LLPS (10, 77), and CHASA also suggests a role for peptide bond exposure in β-turns. The combined surface areas of the C, O and N atoms differ significantly for the central residues in the β-turn ensemble vis a vis the random coil ensemble (**Fig. 3B**), and this may reflect differences in a potential for peptide π/sp^2^ interactions. Other non-turn, multi-turn, multi-valent interactions are equally important to LLPS, but the solvent water considerations, elucidated by CHASA, suggest a plausible reason for why β-turn propensity is elevated in our testing set of LLPS-competent IDRs.

**Figure 3.**
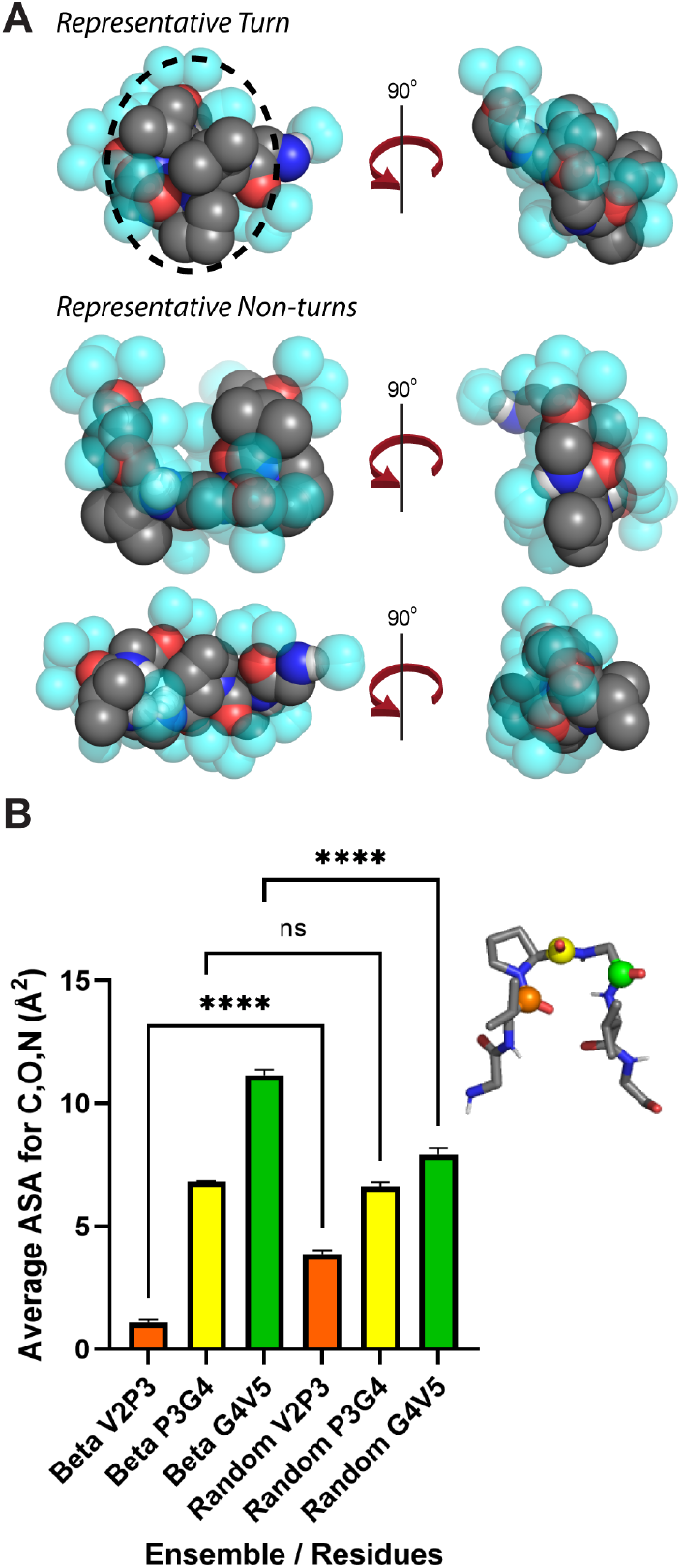
β-turn effects on protein-protein interactions. **A)** Conformations are shown for the ELP repeat GVPGVG, including sterically allowed, CHASA-generated solvation waters (see text). The upper panel shows a representative turn conformation, where hydrophobic accessible surface area is clustered. The lower panels show two representative random coil conformations, and no such clustering is observed. **B)** Surface area for C, O, N atoms in the peptide bond when CHASA waters have been placed for the central turn residues in turn (left) and random coil (right) ensembles. Error bars represent the 95% confidence interval calculated over 1,000 conformations. The significance is calculated using the Welch and Brown-Forsythe one-way ANOVA test for non-equivalent variances with Games-Howell post hoc analysis (**** is p < 0.0001; ns is not significant). The inset demonstrates which peptide bonds are plotted: orange (between V2 and P3), yellow (between P3 and G4), or green (between G4 and V5).

### Sequence calculated internal scaling exponent, *v_int_*, does not identify IDRs from LLPS proteins

When combined with *v_model_*, several amino acid scales of intrinsic β-turn propensity showed the ability to separate protein classes (Fig. 2B-D). Based on this result, we sought to determine if a different sequence-based calculation of *v* likewise could be used to identify IDRs that drive LLPS. We began with previous work using computationally determined *v_int_* values to predict phase separating properties (36). Similar to our work predicting mean *R_h_* from the primary sequence, SAXS data was used to develop a predictor of *v_int_*. The calculation uses sequence hydropathy decoration, *SHD*, and sequence charge decoration, *SCD*,

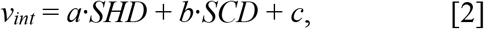

where *a*, *b*, and c are simulation-derived fitting parameters found to be −0.0423, 0.0074, and 0.701, respectively (36). *SHD* is calculated from sequence by *N*^−1^ ∑_*i*_ ∑_*j,j>i*_( *λ*_*i*_ + *λ*_*j*_)|*j-i*|^−1^, where *λ* is the amino acid specific hydropathy (78) normalized to have values from 0 to 1 (36). *SCD* is calculated from sequence by *N*^−1^ ∑_*i*_ ∑_*j,j*>*i*_(*q_i_q_j_*|*j-i*|^½^, where *q* is the amino acid specific charge (79). For the null set, the mean *v_int_* was 0.494 ± 0.083. For the testing set, it was 0.508 ± 0.085. A two-sample z-test comparing *v_int_* in the null and testing sets gave a one-tail p-value of 0.23, providing no evidence for a statistical difference in the means of the two sets. Moreover, sequence calculated *v_int_* and *v_model_* were not correlated when compared (R^2^ = 0.002, Fig. S3). Consistent with the lack of correlation between *v_int_* and *v_modei_*, *v_int_* finds the testing and null sets as statistically similar. As such, these two representations of scaling exponents may exhibit different prediction capabilities for identifying LLPS proteins and protein regions. Sequence patterning is important, and especially hydropathy and charge decoration, but it may not exclusively capture LLPS potential.

### Sequence calculated turn propensity and *v_model_* predict protein regions driving LLPS

Next, we sought to determine if the observed differences in mean *v_model_* and mean β-turn propensity between the null, testing, and folded sets (Fig. 2B) could be used to identify regions within the protein that match the LLPS class, and thus predict the specific protein regions that support a demixing transition. For the initial test, *v_model_* and β-turn propensity were calculated for each Sup35 domain. The sequence of the N-terminal prion domain that mediates phase separation (41) gave 0.531 and 1.183 for *v_model_* and β-turn propensity, respectively, which matched the testing set averages (Fig. 4A). In contrast, sequences representing the ID middle and folded C-terminal domains gave values for *v_model_* and β-turn propensity that were most like the null and folded sets, respectively (Fig. 4A).

**Figure 4.**
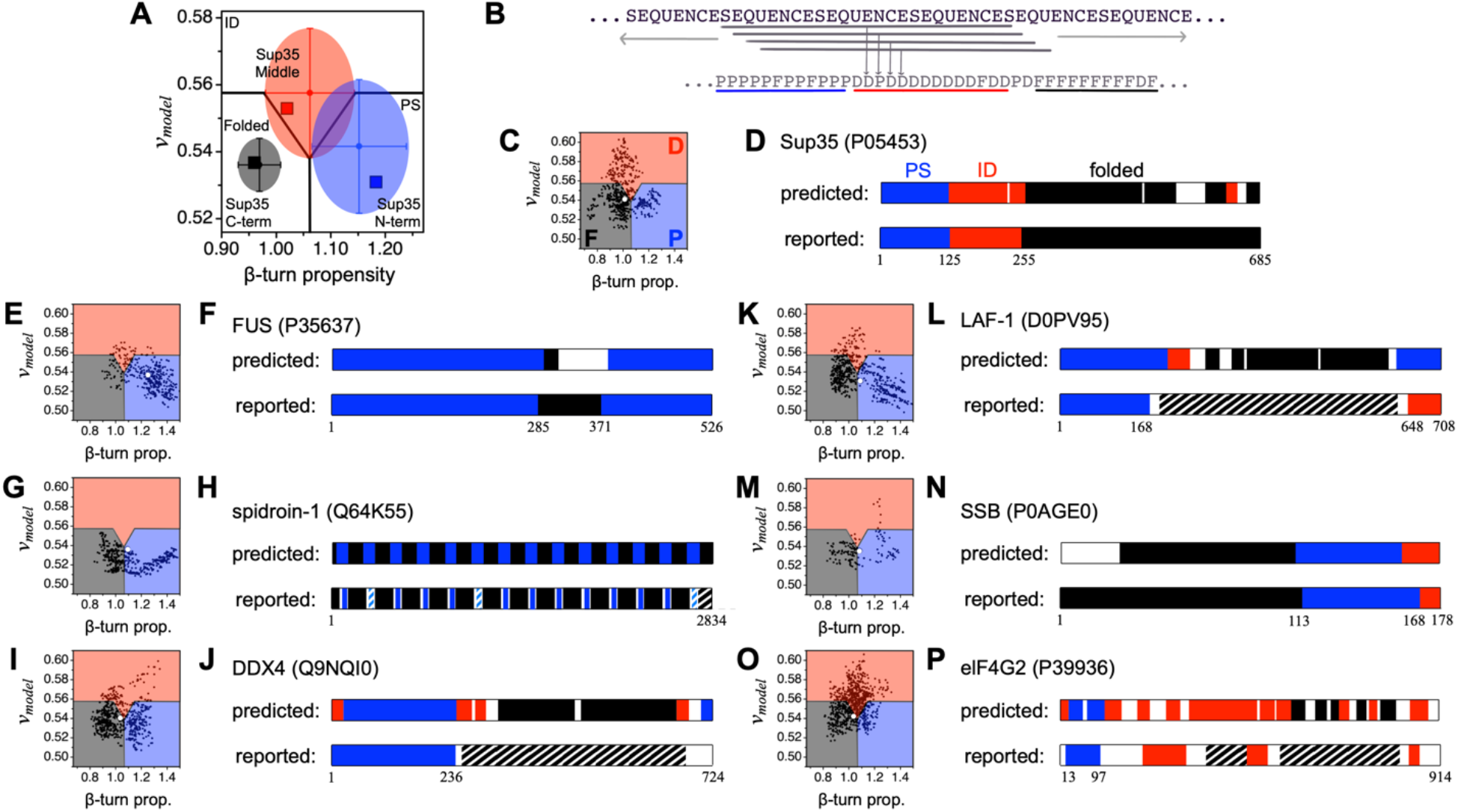
Predicting protein regions that drive LLPS. **A)** Sequence calculated *v_model_* and β-turn propensity for the Sup35 domains – the ID N-terminal domain (blue square), the ID middle domain (red square), and the folded C-terminal domain (black square) – is compared to the set averages (see Fig. 2B). **B)** A sliding window algorithm was used to identify regions within a protein that match the LLPS class. β-turn propensity and *v_model_* are calculated for each contiguous stretch of 25-residues, or “window”, in the primary sequence. **C)** Each window is assigned a label of P, D, or F depending on if *v_model_* and β-turn propensity for the window placed it in the PS, ID, or Folded sector, respectively, of a β-turn propensity vs. *v_model_* plot. This label was given to the central residue of the window. N- and C-terminal residues not belonging to a central window position were assigned the label of the first window and last window, respectively. In this plot, the calculated values of *v_model_* and β-turn for the whole protein sequence is shown by the larger, white dot. **D)** Contiguous regions (*N*≥20) that were 90% of only one label P, D, or F were colored blue, red, or black, respectively, to represent predicted PS, ID, or folded regions. **E-P)** The sliding window calculation was applied to the whole sequences of six additional proteins, identified in the figure by name and UniProt accession number. Striped represents ≥50% identity to a known LLPS IDR (blue) or folded protein (black).

To analyze proteins without using pre-defined boundaries for different regions, we apply a 25-residue window and then slide this window across the whole sequence in 1-residue steps, as shown schematically in Figure 4B. The choice of 25 residues for the window size was arbitrary. For each 25-residue window, *v_model_* and β-turn propensity were calculated from the amino acid sequence of the window. We mapped these values onto a β-turn propensity vs. *v_model_* plot, which was divided into sectors labeled PS, ID, and Folded. Sector boundaries were defined by the mean and standard deviations in *v_model_* and β-turn propensity in the null set (Fig. 4A). To demonstrate this scheme, Figure 4C shows the results from using this algorithm on the full Sup35 sequence, where each small dot in the figure represents a different 25-residue window. The Sup35 primary sequence was then assigned a new three-letter code: P, D, or F based on window localization into the PS, ID, or Folded sectors. We then identified regions in the sequence of length ≥20 residues that were at least 90% of only one of these labels, and color coded those identified regions (Fig. 4D).

We sought to determine if our algorithm would similarly identify regions of proteins with diverse reported mechanisms driving LLPS. Figures 4E-P show the results from applying this algorithm, ParSe, to the whole sequences of six additional proteins that have been characterized *in vitro* and verified to exhibit LLPS. The phase separation of FUS (Fig. 4E, F) under physiological-like protein concentration is known to require the full-length sequence (80). FUS also has a short, folded domain, the RNA recognition motif (RRM) spanning residues 285-371 (81). The silk-wrapping protein, spidroin-1 (Fig. 4G, H), consists of a repeat folded region (82) with intervening, short IDRs that mediate phase separation via hydrophobic amino acids (83). The N terminus (residues 1-236) of DDX4 (Fig. 4I, J) uses a network of charge, hydrophobic, cation-π, and aromatic interactions to drive phase separation, mostly from F and R residues (3, 84). The R- and G-rich N terminus (residues 1-168) of LAF-1 (Fig. 4K, L) is used for both phase separation and binding ssRNA (4), while its core domain (residues 231-628) represents a RecA-like DEAD box helicase containing ATP and RNA binding sites (85). The ID C terminus (residues 648-708) of LAF-1 is not required for phase separation *in vitro* (4) and so may have been incorrectly predicted by the algorithm. LLPS of SSB proteins (Fig. 4M, N) is thought to be driven by the low sequence complexity ID linker region (86) that connects a highly conserved N terminus OB fold (87) to a C-terminal peptide motif. In elF4G2 (Fig. 4O, P), a small N- and Q-rich region (residues 13-97) has been shown to be sufficient for LLPS (88). Also, modelling based on sequence similarity (89) has been used to predict two structured domains in elF4G2, one that was identified by ParSe. For the proteins GRB2 (6) and SPOP (7) that drive phase separation and are mostly folded (90, 91), ParSe did not find any regions with high LLPS potential (Fig. S4). In summary, our algorithm predicted regions driving LLPS in proteins with a variety of reported mechanisms, indicating that *v* and β-turn propensity may represent a unifying property driving LLPS.

We noticed that within eIF4G2, LAF-1 and DDX4 that regions previously annotated as folded contained some sections we predicted to be PS or ID. As our analysis is designed for disordered and not ordered regions, we sought to determine an error rate for misclassifying ordered domains. We took as a larger database of folded domains the set of >14,000 proteins listed by SCOPe (Structural Classification of Proteins extended, version 2.07) that represent the globular fold classes across families and superfamilies (92, 93). Using β-turn propensity and *v_model_* calculated for the full domain of each protein found in SCOPe, we found that 95.4% resided in the folded sector of the β-turn propensity vs. *v_model_* plot. This is comparable to other established ID predictors. For example, using metapredict (94), selected because it can quickly process large numbers of sequences, we found 99.5% of sequences to have average disorder scores less than 0.5 (indicating folded). Of the proteins in the SCOPe database with average disorder score greater than 0.5 by metapredict, only half were also identified as not folded by ParSe, indicating that combining an established ID predictor with our classification scheme could improve its fidelity.

### Long regions with both high β-turn propensity and low *v_model_* are rare in the human proteome

We noticed that most proteins found in the testing set had not just IDRs with a high average β-turn propensity and low average predicted *v_model_*, but that they tended to contain long (≥50 residues) stretches labeled by ParSe to be “P”. To determine whether this feature is unique to proteins driving LLPS, we measured the prevalence of regions predicted from sequence to have high LLPS potential in the human proteome (Fig. 5). These were identified as regions with at least 90% of residue positions labeled as “P” by ParSe (Fig. 4B). We found that ~70% of the human proteome had a region at least 1 residue in length with predicted high LLPS potential (i.e., a single P-labeled position), while only ~4% have such a region that is at least 50 residues in length. This result shows that few human proteins possess a region of substantial length (≥50 residues) that combines high β-turn propensity with low *v_model_*.

**Figure 5.**
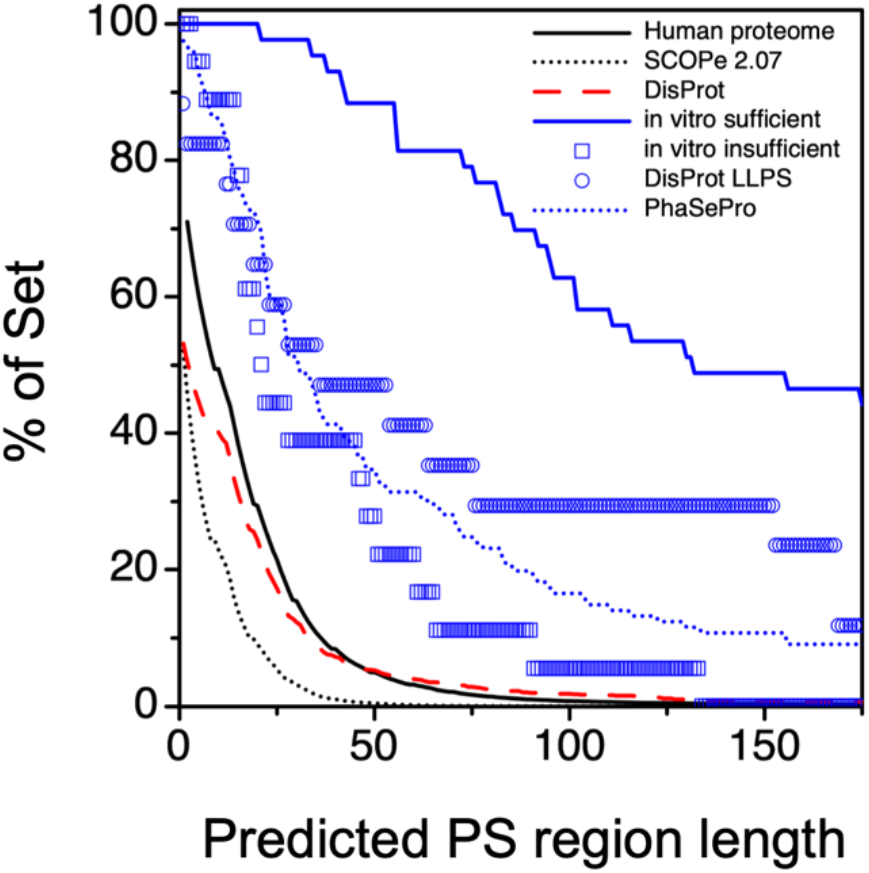
Long regions matching the LLPS IDR class are rarely found in the human proteome, the DisProt database, and folded proteins. The sliding window calculation was used to identify regions in proteins that were ≥90% labeled P (see Fig. 4), which are referred to in this figure as phase separating, PS, regions. Shown by the y-axis is the percent of proteins in a set with a PS region at least as long as the length indicated by the x-axis. The human proteome (UniProt reference proteome UP000005640) is given by the solid, black line; the DisProt database (minus “liquid-liquid phase separation” annotated entries) is given by the dashed red line; and the SCOPe database (version 2.07), representing a wide selection of folded proteins, is given by the black, stippled line. In comparison, long PS regions predicted by ParSe are enriched in the set of *in vitro* sufficient LLPS proteins (solid, blue line), the DisProt “liquid-liquid phase separation” annotated IDPs (open, blue circles), the PhaSePro database (stippled, blue line), and the set of *in vitro* insufficient LLPS proteins (open, blue squares).

Next, we repeated this calculation for the set of 43 proteins assembled by Vernon *et al* (10) that have been verified *in vitro* to exhibit phase separation behavior. Figure 5 shows that almost 90% of these “*in vitro* sufficient” LLPS proteins have a region predicted by our algorithm to have high LLPS potential that is 50 residues in length or longer. Vernon *et al* (10) also prepared a set of 18 additional proteins observed *in cellulo* to exhibit phase separation behavior that were found through *in vitro* characterization not to phase separate as purified proteins (Table S6). Labeled as “*in vitro* insufficient”, less than a third of these proteins (28%) contain a 50-residue or longer region with high LLPS potential, however, ~60% have a 20-residue or longer region with high β-turn propensity and low *v_model_*. The DisProt database, minus the LLPS annotated IDPs, mirrored the human proteome result, demonstrating that ID alone is not sufficient to trigger LLPS prediction by ParSe. The LLPS-annotated IDPs in the DisProt database were enriched in P-labeled regions, giving results in between the *in vitro* sufficient and *in vitro* insufficient Vernon *et al* protein sets, while the PhaSePro database (all proteins) gave results that were most like the *in vitro* insufficient LLPS proteins. The set of proteins in SCOPe were mostly devoid of regions predicted to have high LLPS potential by ParSe (Fig. 5). Thus, while proteins containing long, contiguous P-labeled regions are highly represented in proteins known to undergo LLPS, these regions appear relatively unique to this class of proteins. Supporting this observation, using the Mann-Whitney U test indicates that the distributions of predicted PS region lengths shown in Figure 5 are each significantly different from the others, especially when comparing sets known to be enriched for LLPS to the sets that are not (Table S7).

### β-turn propensity to *v_model_* ratio is correlated with other predictors of phase separation

Having demonstrated that ParSe was able to identify regions driving phase separation, we next sought to determine whether this algorithm was recognizing similar sequence features as other predictors of phase separation (95, 96). Existing predictors are based on molecular mechanisms thought to drive phase separation (10–12), experimental databases of non-specific protein interactions (97), or machine learning outputs based on sequence databases (40). Phase separation may be promoted by many different mechanisms, including β-sheet interactions that also drive prion formation (12), interactions with nucleic acids (11), arginine and tyrosine content (80), and multivalent protein-protein interactions (98). As a result, different predictors will be able to identify different proteins, and any correlation between predictors may indicate an evolutionary relationship between different mechanisms (95, 96).

To facilitate a direct comparison to other predictors, we sought to collapse our predictions to a single value. We noticed that sequences with higher β-turn propensity and lower *v_model_*, and thus larger values of this ratio, were found primarily in the testing set. We hypothesized that *r_model_*, the ratio of β-turn propensity to *v_model_*, would be maximized in IDRs that drive LLPS. Consistent with this idea, mean *r_model_* (± σ) was 2.1 ± 0.2, 1.9 ± 0.1, and 1.8 ± 0.1 in the testing, null and folded sets, respectively. We then compared *r_model_* for the sequences in each set to PScore (10), granule propensity from catGRANULE (11), PSPredict score (40), and LLR from PLAAC (12). We limited our analysis to sequences at least 140 amino acids in length, a PScore requirement, so the same sequences could be compared across all predictors. In addition, we compared *r_model_* calculated from sliding windows to residue-level values provided by PScore and CatGRANULE. The strength of the correlation between all predictors on our testing, null, and folded sets, and their combination, was measured by calculating the coefficient of determination (R^2^, Fig. S5A-F).

Consistent with ParSe’s ability to recognize sequences driving LLPS that utilize a variety of mechanisms (Fig. 4) we found strong correlation between *r_model_* and all four of the predictors, with generally higher R^2^ values than the other pairwise comparisons; granule propensity and PScore are also generally well correlated (Fig. 6). For example, 4 of the 6 pairwise comparisons with R^2^ above 0.5 are with *r_model_*. The pairwise correlation between predictors was, very generally, higher for the testing set than either the null or folded sets. Similarly, the residue-level correlation between *r_model_* and PScore was higher for the testing than the folded and null sets (Fig. S5E). On the other hand, because the testing set typically had high values, whereas the folded set low values, the correlations in the combined set containing all of the training, folded, and null sets, were most often stronger than within each individual set. We speculate that the relatively higher correlation within the testing set than null or folded sets is the result of evolutionary pressure for phase separating IDRs to utilize multiple mechanisms to drive phase separation, e.g., containing both cation-π interactions and a high β-turn propensity within the same sequences. As a first step towards testing this hypothesis, we created a set of 100 random sequences with the same average amino composition as the testing set. The residue level correlation between *r_model_* and PScore was significantly reduced with scrambled sequences, indicating the individual sequences efficiently combine LLPS mechanisms in a way that is not reflected in the amino acid composition averaged across many sequences (Fig. S5E). This is consistent with previous proposals that patterns of self and solvent interactions in a sequence may feature prominently in mechanisms promoting LLPS and for determining condensate specificity (99, 10, 100, 98).

**Figure 6.**
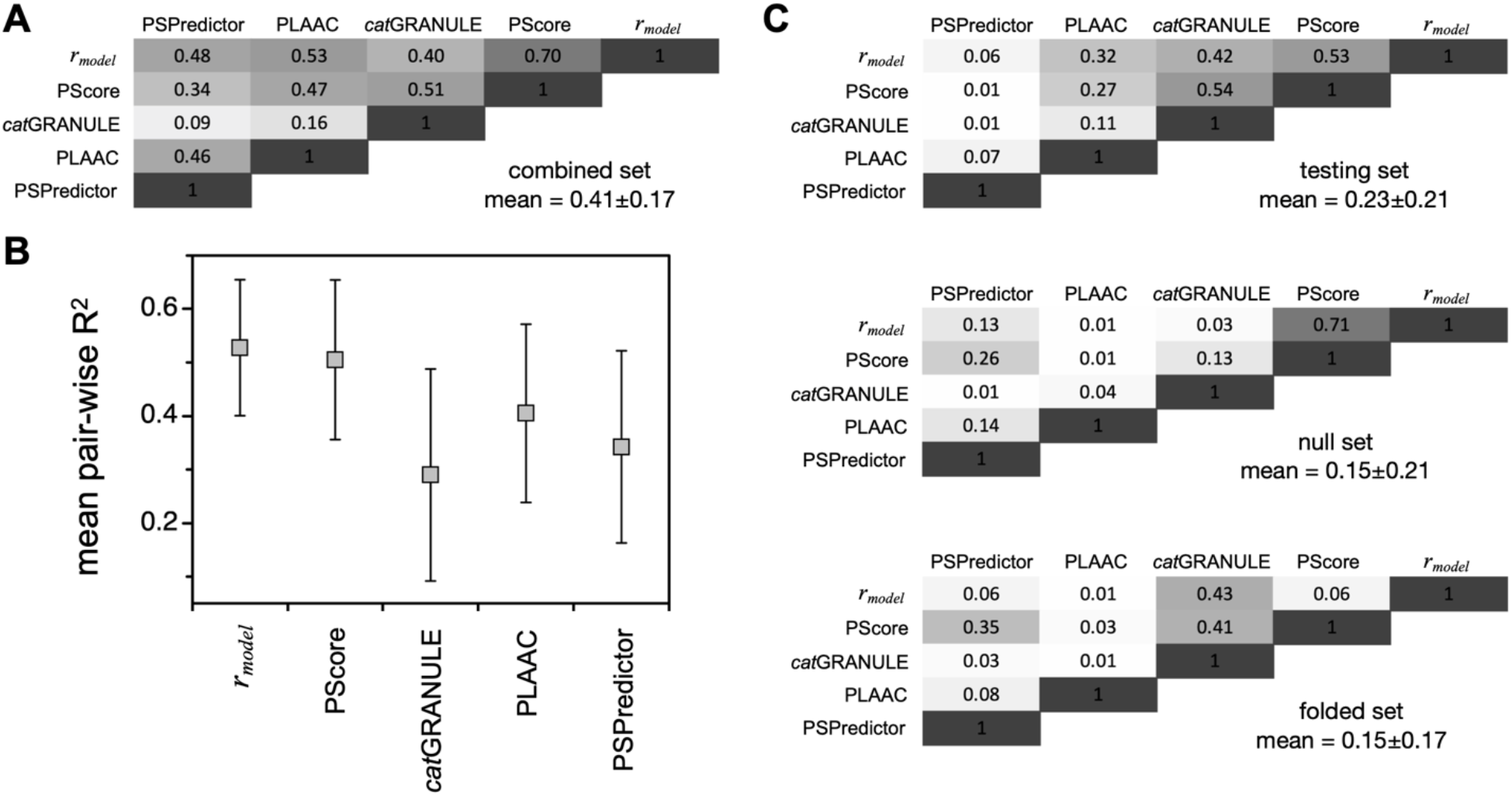
Comparing phase separation predictors. **A)** Pair-wise correlations, R^2^, for *r_model_*, PScore, *cat*GRANULE granule propensity, PLAAC LLR score, and PSPredictor score calculated for the sequences when combining the testing, null, and folded sets. Grayscale indicates the magnitude of R^2^; the mean and standard deviation for all pair-wise combinations in the combined set is shown. **B)** The mean pairwise R^2^ of each predictor, with error bars showing standard deviation, for the combined set correlations (panel A). **C)** Pair-wise correlations, R^2^, calculated for the individual sets: testing (top), null (middle), and folded (bottom). The mean and standard deviation for all pair-wise combinations within a set is shown. Plots of pair-wise correlations involving *r_model_* are in Figure S5.

## Discussion

The hydrodynamic dimensions of proteins have long been studied to investigate folding mechanisms (101), properties of the denatured state ensemble (102), and the physical characteristics of IDPs (3, 13, 19–21, 23). By normalizing hydrodynamic size to the chain length, the predicted polymer scaling exponent, *v*, provides a simple metric that reports on the net balance of self and solvent interactions. This is similar, though not identical to the original intended use of *v* to describe the flexible homopolymer whereby subunit-subunit interactions are all equivalent, as are, separately, subunit-solvent interactions (18). Because IDPs are heteropolymers and contain varying, spatially organized, local interactions, *v* reflects a phenomenological parameterization rather than an exact description of molecular forces present. While there are real limitations to the applicability of applying concepts developed for long homopolymers to heteropolymeric proteins, numerous studies, including this work, support the view that properties, like *v*, derived for homopolymers, can be successfully applied to biological IDPs to help understand their observed solution behavior (19, 24, 25, 31–33).

Here, we have used sequence-based calculations of mean *R_h_*, which has been found to match the measured values from many IDPs (14, 35, 44), to test the wide-spread notion that lower *v* is associated with the potential for LLPS of IDRs (19, 24, 25, 31). Using *v_model_*, obtained from sequence calculated *R_h_*, we find that IDRs from proteins that exhibit LLPS have, on average, reduced *v_model_* when compared to non-phase separating IDRs, but with significant statistical overlap between the two sets (Fig. S2). Thus, it is unlikely that *v_model_*, by itself, has significant predictive power for LLPS. However, β-turn propensity also is different in the different protein classes (Table 2), consistent with the enhancement of the accessibility of π/sp^2^ electronic interactions in β-turns relative to random conformations (Fig. 3). β-turn propensity, when combined with *v_model_*, shows the ability to predict protein regions: for folded, phase separating, and non-phase separating IDRs (Fig. 4). Protein regions having low *v_model_* combined with high β-turn propensity are rare in the human proteome, and especially rare in folded proteins, while enriched in known LLPS proteins (Fig. 5). Because many proteins and peptides can be induced to form phase separated states under different solution conditions (103, 104), we hypothesize that protein regions having low *v_model_* combined with high β-turn propensity identify IDRs that drive phase separation under mostly mild, physiological-like conditions. Other sequence-based predictors of protein LLPS, for example PSCORE (10) and catGRANULE (11), similarly identify only a small subset of the human proteome as exhibiting high LLPS potential. Our work also builds on the recent finding that phase separating IDRs are less hydrophobic by traditional scales than non-phase separating IDRs, yet more compact (105, 106). The compaction appears to occur through other mechanisms than hydrophobicity, including cation-π and charged interactions (10, 80, 106), as well as a high propensity for β-turns (Fig. 4).

To predict protein regions in a given primary sequence, based on calculations of *v_model_* and β-turn propensity, we have written the ParSe algorithm and have made it available online (see Data Availability). In addition to predicting the locations of protein regions, ParSe outputs *r_model_*, the ratio of β-turn propensity to *v_model_*, both for the whole sequence and at the residue level. Using ParSe, we found that proteins that phase separate as purified components have long predicted PS regions, while *in cellulo* observed LLPS proteins that do not phase separate as purified components seem to, in general, have shorter predicted PS regions. When compared, *r_model_* showed strong correlations to other phase separating predictors, and notably those predictors that are based on mechanisms thought to promote LLPS (Fig. 6). As the ParSe model does not directly evaluate for the sequence features proposed by other mechanisms, namely the patterning of either cation-π, π-π or charged amino acids; or nucleic acid binding; the correlation between these metrics points to an evolutionary constraint to include multiple sequence features in regions that promote LLPS (10, 11, 95). More generally, the correlation between predictors that are based on disparate molecular mechanisms will be useful for determining which molecular features are typically combined in LLPS proteins, and which LLPS proteins instead rely on unique molecular grammars (80, 95, 96, 98, 100, 105, 106).

## Experimental procedures

### Protein databases

Lists of proteins that exhibit LLPS behavior were obtained from Vernon *et al* (10), the PhaSePro database (63), and the DisProt database (107), chosen because each contains protein lists that have been curated manually for experimentally verified cases of LLPS. From the Vernon set, we segregated proteins that phase separate *in vitro* as purified components (Table S2) from those that do not (Table S5). From DisProt, protein sequences that phase separate were found by search using the disorder function ontology identifier for liquid-liquid phase separation, IDPO:00041 (64). A set of IDPs not known to phase separate but with monomeric experimental mean *R_h_*, rather than sequence-predicted mean *R_h_*, was assembled from literature reports (35, 44–62). The human proteome reference set UP000005640 (108), the Structural Classification of Proteins – extended (SCOPe) database version 2.07 (92, 93), and the consensus disordered regions from the DisProt database (2021_06) excluding those regions with the ontology identifier for liquid-liquid phase separation, were used as additional negative controls, i.e., sequence lists not enriched for LLPS behavior.

### Mean *R_h_* sequence calculation

The hydrodynamic dimensions of disordered protein ensembles depend strongly on sequence composition. For IDPs, the mean *R_h_* has been shown to be accurately predicted from the intrinsic bias for the polyproline II (PPII) conformation (14, 44) and sequence estimates of the protein net charge (35, 22). The equation to calculate mean *R_h_* for a disordered sequence is

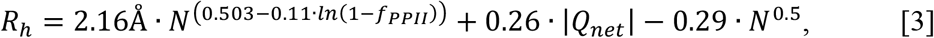

where *N* is the number of residues, *f_PPII_* is the fractional number of residues in the PPII conformation, and *Q_net_* is the net charge (22). *f_PPII_* is estimated from ∑ *P_PPII,i_/N*, where *P_PPII,i_* is the experimental PPII propensity determined for amino acid type *i* in unfolded peptides (109) and the summation is over the protein sequence. *Q_net_* is determined from the number of lysine and arginine residues minus the number of glutamic acid and aspartic acid.

### Disorder prediction

The presence of intrinsic disorder in proteins and protein regions can be predicted from sequence with good confidence (110). The GeneSilico MetaDisorder service (65) was used to calculate the disorder tendency at each position in a sequence from the consensus prediction of 13 primary methods. Residues with a disorder tendency >0.5 are predicted to be disordered, while those with disorder tendency <0.5 are predicted to be ordered. To minimize misidentification, we selected ID regions as those with at least 20 contiguous residue positions having disorder tendency ≥0.7. When the GeneSilico MetaDisorder service was offline or otherwise unavailable, the IUPred2 long predictor (66) was used instead.

### Calculation of β-turn propensity

The propensity to form β-turn structures was calculated by ∑ *scale_i_*/*N*, where *scale_i_* is the value for amino acid type *i* in the normalized frequencies for β-turn from Levitt (37). The summation is over the protein sequence containing *N* number of amino acids. Calculations using the Chou-Fasman normalized β-turn frequencies (38) followed an identical method. Calculations that account for specificity in the four different turn positions (*i*, *i*+1, *i*+2, *i*+3) used the turn potentials from Hutchinson and Thorton (Table 2 in (39)), where a 4-residue window with each residue position in the window a turn position, was slid across the protein sequence in 1-residue increments. For a sequence, the summation of turn potentials in a window was divided by 4, and the overall sum of windows was divided by the number of windows.

### Calculating *v_model_* and β-turn propensity in 25-residue windows for identifying LLPS regions

Our goal was to calculate *v_model_* in a manner that was sensitive to the composition of the window, while also maintaining some independence from the window length that was arbitrarily selected. For example, *v_model_* calculated for all 25-residue windows in the Sup35 primary sequence gives the average value of 0.531. If the window size is doubled by doubling the number of each amino acid type in the window sequence, the average *v_model_* changes to 0.540 despite the same fractional compositions of amino acids in the windows and the same number of windows. Owing to the second and third terms in equation [3], identical fractional compositions of amino acids can yield different *v_model_* depending on window length. To avoid this, we calculated the average *v_model_* for all 25-residue windows in Sup35 at multiples of 1x, 2x, 3x, etc., where “1x” means the amino acid distributions are identical to the native sequence, “2x” doubles the occurrence of each amino acid type in a window, “3x” triples the occurrence, and so forth. By this scheme, the fractional ratio of each amino acid type in a window is constant. We found that the average calculated *v_model_* for 25-residue windows stabilized for multiples ≥4x. Specifically, for fractional compositions obtained from a biological sequence, *v_model_* became length-independent for *N*≥100 while also remaining highly sensitive to changes in the fractional composition. Based on this finding, *v_model_* for a 25-residue window was calculated from sequence by first multiplying the number of each amino acid type in the window sequence by 4 and then by calculating *v_model_* for the resulting 100-residue length. The β-turn propensity for a 25-residue window was calculated without modification to the method as described above and used the normalized turn frequencies from Levitt (37).

### ParSe calculation

For an input primary sequence, whereby the amino acids are restricted to the 20 common types, ParSe first reads the sequence to determine its length, *N*, and the number of each amino acid type. Using these sequence-defined values, *Q_net_* and *f_PPII_* are calculated (described above) and used with *N* to determine *R_h_* by equation [3], which in turn is used to determine *v_model_* by equation [1]. β-turn propensity is calculated as the sequence sum divided by *N* (described above) from the normalized frequencies by Levitt (37). *r_model_* is determined by the ratio of β-turn propensity to *v_model_*. Next, ParSe uses a sliding window scheme (Fig. 4B) to calculate *v_model_* and β-turn propensity for every 25-residue segment of the primary sequence (described above). This window scheme can be applied to proteins with *N* >25. The values of *v_model_* and β-turn propensity calculated for a window determine the window’s localization to a PS, ID, or Folded sector in a β-turn propensity vs. *v_model_* plot (Fig. 4C). The sector boundaries are shown in Figure 4A, and these boundaries are defined by the mean and standard deviation in β-turn propensity and *v_model_* calculated in the null set (Tables 1 and 2). If a window, based on its β-turn propensity and *v_model_* values, is localized to the PS sector, the central residue in that window is labeled “P”, whereas localization to the ID sector labels the central residue position “D”, and localization to the Folded sector labels the residue “F”. N- and C-terminal residues not belonging to a central window position are assigned the label of the central residue in the first and last window, respectively, of the whole sequence. Protein regions predicted by ParSe to be PS, ID, or Folded are determined by finding contiguous residue positions of length ≥20 that are ≥90% of only one label P, D, or F, respectively. When overlap occurs between adjacent predicted regions, owing to the up to 10% label mixing allowed, this overlap is split evenly between the two adjacent regions.

### PSCORE calculation

PSCORE, which is a phase separation propensity predictor (10), was calculated by computer algorithm using the Python script and associated database files available at https://doi.org/10.7554/eLife.31486.022.

### Granule propensity calculation

Granule propensity was calculated by using the *cat*GRANULE (11) webtool available at http://www.tartaglialab.com.

### PLAAC LLR calculation

LLR score, which identifies prion-containing sequences (12), was calculated by using the webtool available at http://plaac.wi.mit.edu.

### PSPredictor calculation

PSPredictor score, which predicts phase separation potential (40), was calculated by using the webtool available at http://www.pkumdl.cn:8000/PSPredictor.

### Metapredict calculation

Metapredict score (94), which predicts the presence of ID in a sequence, was calculated by computer algorithm using the Python script available at http://metapredict.net.

### Computer generation of disordered ensembles

Structures of GVPGVG were generated by a random search of conformational space using a hard sphere collision model (111). This model uses van der Waals atomic radii (112, 113) as the only scoring function to eliminate grossly improbable conformations. The procedure to generate a random conformer starts with a unit peptide and all other atoms for a chain are determined by the rotational matrix (114). Backbone atoms are generated from the dihedral angles φ, ψ, and ω and the standard bond angles and bond lengths (115). Backbone dihedral angles are assigned randomly, using a random number generator based on Knuth’s subtractive method (116). (φ, ψ) is restricted to the allowed Ramachandran regions (117) to sample conformational space efficiently. For peptide bonds, ω had a Gaussian fluctuation of ± 5% about the *trans* form (180°) for nonproline residues. Proline sampled the *cis* form (0°) at a rate of 10% (118). Of the two possible positions of the Cβ atom in nonglycine residues, the one corresponding to L-amino acids was used. The positions of all other side chain atoms were determined from random sampling of rotamer libraries (119). Structures adopting the type II β-turn were identified as those with (φ, ψ) angles of (−60°±15°, 120°±15°) and (80°±15°, 0°±15°) for P3 and G4, respectively, while also containing a hydrogen bond connecting the carbonyl oxygen of V2 to the amide proton of V5. Structures were generated until we had 1,000 turn and 1,000 non-turn structures of the peptide GVPGVG. A variety of structural measurements were taken on each ensemble, and statistical convergence was confirmed by comparing the average values of the first 500 structures to the average over the entire ensemble. Specifically, the average total accessible surface area, end to end distance, and radius of gyration for the first 500 structures was found to be within one standard deviation of the average over the entire ensemble, suggesting that additional conformations did not alter the measurements beyond the first 500 structures.

### CHASA analysis and molecular docking

Computer generated structures, described above, were processed using the CHASA module (76) of the LINUS software package (120, 121). Two structures containing turns were docked using the GOLD/HERMES molecular docking software version 2020.1 (122). After hydrogen atoms were added, docking used the ChemPLP scoring function. The beta carbon on the third proline residue defined the binding site. Valine side chains were sampled using the built-in rotamer library, and all backbone torsions were held fixed in their original conformation. HERMES was used to calculate the buried hydrophobic accessible surface area upon formation of the complex.

## Data availability

Source code. The ParSe algorithm written in Fortran, Parse.f, can be downloaded at https://github.com/stevewhitten/Parse, DOI: 10.5281/zenodo.5138428. A webtool version can be used at http://folding.chemistry.msstate.edu/utils/parse.html.

## Supporting information

This article contains supporting information.

## Author contributions

E.A.P., J.H.A., N.C.F., L.E.H., and S.T.W. performed research. E.A.P., J.J.C., N.C.F., L.E.H., and S.T.W. analyzed data. L.E.H. and S. T. W. conceived the idea. N.C.F., L.E.H, and S.T.W. designed research. N.C.F., L.E.H, and S.T.W. wrote the manuscript. E.A.P., J.H.A., and J.J.C. edited the manuscript.

## Funding and additional information

This work was supported by the National Institutes of Health under grants R15GM115603 (S.T.W.), R25GM102783 (South Texas Doctoral Bridge Program; N. M. J. Blake, B. O. Oyajobi, and S.T.W.), R35GM119755 (L.E.H.), and R01AI139479 (N.C.F.), as well as the National Science Foundation under grants 1818090 (N.C.F.) and 1943488 (L.E.H). No nongovernmental sources were used to fund this project. The content is solely the responsibility of the authors and does not necessarily represent the official views of the NSF or NIH.

## Abbreviations

ID: intrinsically disordered
IDP: intrinsically disordered protein
IDR: intrinsically disordered region
LLPS: liquid-liquid phase separation
ELP: elastin-like polypeptide
CHASA: conditional hydrophobic accessible surface area
ASA: accessible surface area
SAXS: small-angle x-ray scattering
SHD: sequence hydropathy decoration
SCD: sequence charge decoration
PS: phase separating
SCOPe: structural classification of proteins extended.

