## Supporting Information for "Beta turn propensity and a model polymer scaling exponent identify disordered proteins that phase separate"

### Contents:

Supporting Tables
S1. List of IDPs with experimental mean $R_h$.
S2. List of proteins that exhibit phase separation behavior.
S3. Summary of Mann-Whitney U tests that compare mean $v_{model}$ (top) and mean β-turn propensity (bottom) in the null, testing, and folded sets.
S4. Normalized frequency for β-turn.
S5. Structural properties of turn and non-turn ensembles.
S6. List of proteins that exhibit phase separation behavior *in cellulo* that were found by *in vitro* characterization not to phase separate as purified proteins.
S7. Summary of pair-wise Mann-Whitney U tests comparing the relative population (given by set percentage) of predicted PS region lengths, for lengths ranging from 1 to 150 residues.

Supporting Figures
S1. Experimental mean $R_h$ compared to sequence calculated mean $R_h$.
S2. Distribution of $v_{model}$ values in the A) testing and B) null sets.
S3. Comparing sequence calculated $v_{model}$ and $v_{int}$.
S4. Predicting protein regions that drive LLPS.
S5. Pair-wise correlations of predictor results.

Supporting References

### Supporting Tables

#### Table S1. List of IDPs with experimental mean $R_h$.

| Name | $N$ | mean $R_h$ (Å) | sequence | reference |
|---|---|---|---|---|
| p53(1-93) | 95 | 32.4 | GSMEEPQSDPSVEPPLSQETFSDLWKLLPENNVLSPLPSQ<br>AMDDLMLSPDDIEQWFTEDPGPDEAPRMPEAAPPVAPA<br>PAAPTPAAPAPAPSWPL | (1) |
| p53(1-73) TAD | 73 | 23.8 | MEEPQSDPSVEPPLSQETFSDLWKLLPENNVLSPLPSQA<br>MDDLMLSPDDIEQWFTEDPGPDEAPRMPEAAPRV | (2) |
| Vmw65 | 89 | 28 | GSAGHTRRLSTAPPTDVSLGDELHLDGEDVAMAHADAL<br>DDFDLDMLGDGDSPGPGFTPHDSAPYGALDMADFEFEQ<br>MFTDALGIDEYGG | (3) |
| Hdm2-ABD | 97 | 31.7 | ERSSSSESTGTPSNPDLDAGVSEHSGDWLDQDSVSDQFS<br>VEFEVESLDSEDYSLSEEGQELSDEDDEVYQVTVYQAGE<br>SDTDSFEEDPEISLADYWK | (4) |
| prothymosin-α | 110 | 33.6 | MSDAAVDTSSEITTKDLKEKKEVVEEAENGRDAPANGN<br>ANEENGEQEADNEVDEEEEEGGEEEEEEEEGDGEEEDG<br>DEDEEAESATGKRAAEDDEDDDVDTKKQKTDEDD | (4) |
| HIF1-α-403 | 202 | 44.3 | PAAGDTIISLDFGSNDTETDDQQLEEVPLYNDVMLPSPN<br>EKLQNINLAMSPLPTAETPKPLRSSADPALNQEVALKLE<br>PNPESLELSFTMPQIQDQTPSPSDGSTRQSSPEPNSPSEYC<br>FYVDSDMVNEFKLELVEKLFAEDTEAKNPFSTQDTDLD<br>LEMLAPYIPMDDDFQLRSFDQLSPLESSSASPESASPQST<br>VTVFQ | (5) |
| Fos-AD | 168 | 35 | GSHMSVASLDLTGGLPEVATPESEEAFTLPLLNDPEPKPS<br>VEPVKSISSMELKTEPFDDFLFPASSRPSGSETARSVPDM<br>DLSGSFYAADWEPLHSGSLGMGPMATELEPLCTPVVTC<br>TPSCTAYTSSFVFTYPEADSFPSCAAAHRKGSSSNEPSSD<br>SLSSPTLLAL | (6) |
| Mlph(147-240) | 97 | 28 | RLQGGGGSEPSLEEGNGDSEQTDEDGDLDTEARDQPLN<br>SKKKKRLLSFRDVDFEEDSDHLVQPCSQTLGLSSVPESA<br>HSLQSLSGEPYSEDTTSLEP | (7) |
| tau-K45 | 198 | 45 | MSSPGSPGTPGSRSRTPSLPTPPTREPKKVAVVRTPPKSPS<br>SAKSRLQTAPVPMPDLKNVKSKIGSTENLKHQPGGGKV<br>QIINKKLDLSNVQSKCGSKDNIKHVPGGGSVQIVYKPVD<br>LSKVTSKCGSLGNIHHKPGGGQVEVKSEKLDFKDRVQS<br>KIGSLDNITHVPGGGNKKIETHKLTFRENAKAKTDHGAE<br>IVY | (8) |
| Mlph(147-403) | 260 | 49 | RLQGGGGSEPSLEEGNGDSEQTDEDGDLDTEARDQPLN<br>SKKKKRLLSFRDVDFEEDSDHLVQPCSQTLGLSSVPESA<br>HSLQSLSGEPYSEDTTSLEPEGLEETGARALGCRPSPEVQ<br>PCSPLPSGEDAHAELDSPAASCKSAFGTTAMPGTDDVRG<br>KHLPSQYLADVDTSDEDSIQGPRAASQHSKRRARTVPET<br>QILELNKRMSAVEHLLVHLENTVLPPSAQEPTVETHPSA<br>DTEEETLRRRLEELTSNISGSSTSSE | (7) |
| p57-ID | 73 | 24 | VRTSACRSLFGPVDHEELSRELQARLAELNAEDQNRWD<br>YDFQQDMPLRGPGRLQWTEVDSDSVPAFYRETVQV | (9) |
| PDE-γ | 87 | 24.8 | MNLEPPKAEIRSATRVMGGPVTPRKGPPKFKQRQTRQF<br>KSKPPKKGVQGFGDDIPGMEGLGTDITVICPWEAFNHLE<br>LHELAQYGII | (10) |
| LJIDP1 | 94 | 24.52 | MARSFTNIKAISALVAEEFSNSLARRGYAATAQSAGRVG<br>ASMSGKMGSTKSGEEKAAAREKVSWVPDPVTGYYKPE<br>NIKEIDVAELRSAVLGKN | (11) |
| cad136 | 136 | 28.1 | RLEQYTSAVVGNKAAKPAKPAASDLPVPAEGVRNIKSM<br>WEKGNVFSSPGGTGTPNKETAGLKVGVSSRINEWLTKT<br>PEGNKSPAPKPSDLRPGDVSGKRNLWEKQSVEKPAASSS<br>KVTATGKKSETNGLRQFEKEP | (12) |
| α-synuclein | 140 | 28.2 | MDVFMKGLSKAKEGVVAAAEKTKQGVAEAAGKTKEG<br>VLYVGSKTKEGVVHGVATVAEKTKEQVTNVGGAVVTG<br>VTAVAQKTVEGAGSIAAATGFVKKDQLGKNEEGAPQE<br>GILEDMPVDPDNEAYEMPSEEGYQDYEPEA | (13) |
| CFTR-R-region | 189 | 32 | GAMESAERRNSILTETLHRFSLEGDAPVSWTETKKQSFK<br>QTGEFGEKRKNSILNPINSIRKFSIVQKTPLQMNGIEEDSD<br>EPLERRLSLVPDSEQGEAILPRISVISTGPTLQARRRQSVL | (14) |

| | | | | |
|---|---|---|---|---|
| | | | NLMTHSVNQGQNIHRKTTASTRKVSLAPQANLTELDIYS RRLSQETGLEISEEINEEDLKECLFDDME | |
| SNAP25 | 206 | 39.7 | MAEDADMRNELEEMQRRADQLADESLESTRRMLQLVE ESKDAGIRTLVMLDEQGEQLERIEEGMDQINKDMKEAE KNLTDLGKFCGLCVCPCNKLKSSDAYKKAWGNNQDGV VASQPARVVDEREQMAISGGFIRRVTNDARENEMDENL EQVSGIIGNLRHMALDMGNEIDTQNRQIDRIMEKADSNK TRIDEANQRATKMLGSG | (15) |
| ShB-C | 146 | 32.9 | MTLGQHMKKSSLSESSSDMMDLDDGVESTPGLTETHPG RSAVAPFLGAQQQQQQPVASSLSMSIDKQLQHPLQQLT QTQLYQQQQQQQQQQQNGFKQQQQQTQQQLQQQQSH TINASAAAATSGSGSSGLTMRHNNALAVSIETDV | (16) |
| HIF1-α-530 | 170 | 38.3 | NEFKLELVEKLFAEDTEAKNPFSTQDTDLDLEMLAPYIP MDDDFQLRSFDQLSPLESSSASPESASPQSTVTVFQQTQI QEPTANATTTTATTDELKTVTKDRMEDIKILIASPSPTHI HKETTSATSSPYRDTQSRTASPNRAGKGVIEQTEKSHPRS PNVLSVALSQR | (5) |
| Securin | 202 | 39.7 | MATLIYVDKENGEPGTRVVAKDGLKLGSGPSIKALDGR SQVSTPRFGKTFDAPPALPKATRKALGTVNRATEKSVKT KGPLKQKQPSFSAKKMTEKTVKAKSSVPASDDAYPEIEK FFPFNPLDFESFDLPEEHQIAHLPLSGVPLMILDEERELEK LFQLGPPSPVKMPSPPWESNLLQSPSSILSTLDVELPPVCC DIDI | (17) |
| sml1 | 104 | 23.4 | MQNSQDYFYAQNRCQQQQAPSTLRTVTMAEFRRVPLPP MAEVPMLSTQNSMGSSASASASSLEMWEKDLEERLNSI DHDMNNNKFGSGELKSMFNQGKVEEMDF | (18) |
| PGR | 135 | 37.7 | AEPGKPAEPGKPAEPGKPAEPGTPAEPGKPAEPGTPAEP GKPAEPGKPAEPGKPAEPGKPAEPGTPAEPGTPAEPGKP AEPGTPAEPGKPAEPGTPAEPGKPAESGKPVEPGTPAQS GAPEQPNRSMHSTDNKNQ | (4) |
| Aβ(1-40) | 40 | 14.36 | DAEFRHDSGYEVHHQKLVFFAEDVGSNKGAIIGLMVGG VV | (19) |

**Table S2. List of proteins that exhibit phase separation behavior.**

| Name | Database [a] | UniProt accession number | ID regions (*N*) [b] | folded regions (*N*) [c] | PDB entries |
|---|---|---|---|---|---|
| TAF15 | Vernon *et al* | Q92804 | 1-36 (36)<br>62-207 (146)<br>210-234 (25)<br>379-398 (20) | 231-323 (93) | 2mmy.pdb |
| ROA1 | Vernon *et al* | P09651-2 | 183-216 (34)<br>297-320 (24) | 8-181 (174) | 1l3k.pdb |
| laf1 | Vernon *et al* | D0PV95 | 1-194 (194)<br>629-708 (80) | none | N/A |
| FUS | Vernon *et al* | H3BNZ4 | 1-263 (263) | none | N/A |
| DDX3X | Vernon *et al* | O00571 | 20-133 (114)<br>581-637 (57) | 134-580 (447) | 5e7i.pdb<br>2i4i.pdb |
| DDX4 | Vernon *et al* | Q9NQI0 | 26-250 (225)<br>694-724 (31) | none | N/A |
| eIF4H | Vernon *et al* | Q15056 | 10-39 (30)<br>117-152 (36)<br>159-248 (90) | none | N/A |
| NSP1 | Vernon *et al* | P14907 | 1-629 (629) | none | N/A |
| EWS | Vernon *et al* | F8WC90 | 126-164 (39)<br>177-292 (116) | none | N/A |
| TIA1 | Vernon *et al* | P31483 | 338-386 (49) | 2-39 (38)<br>44-85 (42)<br>105-285 (181) | 6eld.pdb<br>2mjn.pdb |
| Elastin | Vernon *et al* | P15502 | 466-488 (23)<br>611-651 (41) | none | N/A |
| ROA2 | Vernon *et al* | P22626-2 | 184-221 (38) | 3-183 (181) | 5en1.pdb |
| fib1 | Vernon *et al* | P22232 | 51-77 (27) | none | N/A |
| pgl-3 | Vernon *et al* | G5EBV6 | 445-466 (22)<br>519-589 (71)<br>616-693 (78) | none | N/A |
| CIRBP | Vernon *et al* | Q14011 | 86-172 (87) | 3-85 (83) | 5tbx.pdb |
| thermoNup98 | Vernon *et al* | D3KYQ3 | 88-120 (33)<br>125-167 (43) | none | N/A |
| tbNup158 | Vernon *et al* | Q387F2 | 1-50 (50)<br>120-147 (28) | none | N/A |
| Silk (spidroin-1) | Vernon *et al* | Q64K55 | 77-96 (20) [e] | 122-259 (138) [f] | 2mu3.pdb |
| Nup100 | Vernon *et al* | Q02629 | 20-49 (30)<br>215-238 (24)<br>357-384 (28)<br>416-438 (23)<br>445-465 (21)<br>763-789 (27) | none | N/A |
| Nup116 | Vernon *et al* | Q02630 | 259-279 (21)<br>475-527 (53)<br>904-942 (39) | 967-1111 (145) | 3pbp.pdb |

| | | | | | |
|---|---|---|---|---|---|
| Nup98B | Vernon *et al* | F4ID16 | 373-406 (34)<br>421-459 (39)<br>767-836 (70) | none | N/A |
| ddNup220 | Vernon *et al* | Q54EQ8 | 89-130 (42)<br>284-341 (58)<br>358-432 (75)<br>437-520 (84)<br>534-559 (26)<br>887-925 (39)<br>928-1017 (90)<br>1175-1198 (24) | none | N/A |
| TDP43 | Vernon *et al* | Q13148 | 344-371 (28) | 2-79 (78)<br>103-180 (78)<br>320-343 (24) | 5mdi.pdb<br>4y0f.pdb<br>2n3x.pdb |
| xNup214 | Vernon *et al* | Q9PVZ2 | 987-1032 (46)<br>1193-1213 (21)<br>1479-1503 (25) | none | N/A |
| ceNup98 | Vernon *et al* | G5EEH9 | 715-737 (24) | none | N/A |
| Nup98 | Vernon *et al* | P52948 | 521-565 (45)<br>886-940 (55) | 158-213 (56)<br>729-887 (159) | 3mmy.pdb<br>1ko6.pdb<br>2q5x.pdb |
| Nup153 | Vernon *et al* | P49790 | 98-123 (27)<br>404-424 (21)<br>1316-1340 (25)<br>1360-1386 (27) | none | N/A |
| xNup58 | Vernon *et al* | Q5EAX5 | none | 283-406 (124) | 5c3l.pdb |
| m. BugZ | Vernon *et al* | Q9JMD0-3 | 91-185 (95)<br>206-464 (259) | none | N/A |
| bfNup98 | Vernon *et al* | C3XWA2 | 636-677 (42)<br>691-716 (26)<br>876-979 (104)<br>1016-1039 (24) | none | N/A |
| xNup98 | Vernon *et al* | J7I6Y1 | 618-667 (50)<br>877-926 (93)<br>938-960 (23) | 716-866 (151) | 5e0q.pdb |
| dmNup98 | Vernon *et al* | Q9VCH5 | 703-802 (100) | none | N/A |
| xNup153 | Vernon *et al* | K9ZRR1 | 394-418 (25) | none | N/A |
| RBM14 | Vernon *et al* | Q96PK6 | none | 77-153 (77) | 2dnp.pdb |
| xNup54 | Vernon *et al* | K9ZTJ6 | none | 214-450 (237) | 5c2u.pdb<br>5c3l.pdb |
| x. BugZ | Vernon *et al* | Q7ZXV8 | 91-156 (66)<br>185-224 (40)<br>277-323 (47)<br>326-409 (84)<br>413-445 (33) | none | N/A |
| xNup62 | Vernon *et al* | Q91349 | none | 358-485 (128) | 5c3l.pdb |
| xPom121 | Vernon *et al* | Q5EWX9 | 439-466 (28) | none | N/A |
| WHI3 | Vernon *et al* | P34761 | 244-280 (37)<br>379-413 (35) | none | N/A |

| | | | | | |
|---|---|---|---|---|---|
| | | | 446-503 (62)<br>524-543 (20) | | |
| x1CG1 | Vernon *et al* | Q5XGN1 | none | none | N/A |
| xNup50 | Vernon *et al* | Q6DEC7 | 287-320 (34) | none | N/A |
| meg-3 | Vernon *et al* | Q9TXM1 | 1-38 (38)<br>84-105 (22)<br>150-177 (28)<br>271-295 (25)<br>473-493 (21)<br>508-536 (29) | none | N/A |
| eIF4G2 | Vernon *et al* | P39936 | 1-85 (85)<br>127-154 (28)<br>163-225 (63)<br>240-298 (59)<br>308-339 (32)<br>479-522 (44)<br>823-914 (92) | none | N/A |
| FMRP | PhaSePro | Q06787 | 434-632 (199) | 2-207 (206)<br>219-425 (207) | 4qvz.pdb<br>2qnd.pdb |
| TDP43 | PhaSePro | Q13148 | 270-306 (37)<br>350-374 (25)<br>384-414 (31) | 2-79 (78)<br>102-269 (168)<br>307-349 (43) | 5mdi.pdb<br>4bs2.pdb<br>2n2c.pdb |
| Nephrin | PhaSePro | O60500 | 478-502 (25)<br>1024-1061 (38)<br>1095-1229 (135) | none | N/A |
| N-WASP | PhaSePro | O00401 | 140-163 (24)<br>174-206 (33)<br>271-505 (235) | 207-270 (64) | 2lnh.pdb |
| NCK1 | PhaSePro | P16333 | 252-271 (20) | 4-59 (56)<br>101-163 (63)<br>281-377 (97) | 5qu2.pdb<br>2cub.pdb<br>2ci8.pdb |
| NUP98 | PhaSePro | P52948 | 39-81 (43)<br>103-157 (55)<br>214-241 (28)<br>254-346 (93)<br>375-434 (60)<br>447-591 (145)<br>607-686 (80)<br>698-728 (31)<br>883-982 (100)<br>995-1038 (44)<br>1069-1120 (51) | 158-213 (56)<br>729-880 (152) | 3mmy.pdb<br>2q5y.pdb |
| HP1α | PhaSePro | P45973 | 72-110 (39) | 17-68 (52)<br>111-170 (60) | 3fdt.pdb<br>3i3c.pdb |
| NPM1 | PhaSePro | P06748 | 120-239 (120) | 14-119 (106)<br>240-294 (55) | 5ehd.pdb<br>2llh.pdb |
| UBQLN2 | PhaSePro | Q9UHD9 | 104-143 (40)<br>167-189 (23)<br>200-353 (154)<br>369-473 (104)<br>483-624 (142) | 1-103 (103) | 1j8c.pdb |
| HSPB2 | PhaSePro | Q16082 | 1-27 (27)<br>162-182 (21) | 70-149 (80) | 6f2r.pdb,<br>chain c |

| | | | | | |
|---|---|---|---|---|---|
| MAPT | PhaSePro | P10636-8 | 1-343 (343)<br>356-430 (75) | none | N/A |
| TNRC6B | PhaSePro | Q9UPQ9 | 1-1269 (1269)<br>1295-1638 (344)<br>1724-1764 (41)<br>1775-1833 (59) | none | N/A |
| Galectin-3 | PhaSePro | P17931 | 1-105 (105) | 113-250 (138) | 1kjl.pdb |
| p62 | PhaSePro | Q13501 | 194-388 (195) | 4-26 (23)<br>41-89 (49)<br>126-169 (44)<br>389-436 (48) | 6jm4.pdb<br>6jm4.pdb<br>5yp7.pd<br>1q02.pdb |
| LAT | PhaSePro | O43561 | 61-121 (61)<br>136-262 (127) | none | N/A |
| GRB2 | PhaSePro | P62993 | none | 1-217 (217) | 1gri.pdb |
| SOS1 | PhaSePro | Q07889 | 1047-1333 (287) | 6-1046 (1041) | 3ksy.pdb<br>1nvv.pdb |
| NONO | PhaSePro | Q15233 | 1-65 (65)<br>305-471 (167) | 66-304 (239) | 3sde.pdb |
| SFPQ | PhaSePro | P23246 | 1-278 (278)<br>598-707 (110) | 279-597 (319) | 6ncq.pdb<br>4wik.pdb |
| SYN1 | PhaSePro | P17600 | 1-115 (115)<br>380-705 (326) | none | N/A |
| cGAS | PhaSePro | Q8N884 | 1-153 (153) | 154-522 (369) | 4lev.pdb |
| MED1 | PhaSePro | Q15648 | 535-1581 (1047) | none | N/A |
| BRD4 | PhaSePro | O60885 | 1-43 (43)<br>173-346 (174)<br>464-600 (137)<br>684-1351 (668) | 44-172 (129)<br>347-463 (117)<br>601-683 (83) | 5u2e.pdb<br>6ffd.pdb<br>6bnh.pdb |
| Sam68 | PhaSePro | Q07666 | 1-98 (98)<br>183-219 (37)<br>274-440 (167) | 99-135 (37) | 2xa6.pdb |
| HNRNPD | PhaSePro | Q14103 | 1-95 (95)<br>267-290 (24) | 98-175 (78)<br>181-259 (79) | 1hd0.pdb<br>5im0.pdb<br>1wtb.pdb |
| SPOP | PhaSePro | O43791 | none | 28-356 (329) | 3hqi.pdb<br>4j8z.pdb |
| DAXX | PhaSePro | Q9UER7 | 5-54 (50)<br>143-182 (40)<br>387-740 (354) | 55-140 (86)<br>183-386 (204) | 5y18.pdb<br>4h9n.pdb |
| RPB1 | PhaSePro | P24928 | 36-60 (25)<br>322-355 (34)<br>605-624 (20)<br>718-759 (42)<br>1500-1970 (471) | none | N/A |
| Cyclin-T1 | PhaSePro | O60563 | 266-286 (21)<br>297-726 (430) | 4-263 (260) | 3blh.pdb<br>2pk2.pdb |
| DYRK1A | PhaSePro | Q13627 | 27-89 (63)<br>114-133 (20)<br>482-762 (281) | 134-481 (348) | 2vx3.pdb |
| PrP | PhaSePro | P04156 | 22-115 (94) | 119-230 (112) | 1i4m.pdb<br>1fkc.pdb |

| | | | | | |
|---|---|---|---|---|---|
| CBX2 | PhaSePro | Q14781 | 63-275 (213)<br>286-516 (231) | 9-62 (54) | 5epk.pdb |
| TIS11B | PhaSePro | Q07352 | 44-112 (69)<br>217-251 (35)<br>263-329 (67) | none | N/A |
| GATA3 | PhaSePro | P23771 | 2-223 (222)<br>235-260 (26)<br>366-443 (78) | 261-365 (105) | 4hc9.pdb |
| ERα | PhaSePro | P03372 | 104-174 (71)<br>256-281 (26)<br>551-572 (22) | 180-252 (73)<br>305-548 (244) | 1hcp.pdb<br>2ocf.pdb |
| DYRK3 | PhaSePro | O43781 | 1-35 (35)<br>45-137 (93) | 138-532 (395) | 5y86.pdb |
| SYN2 | PhaSePro | Q92777 | 1-115 (115)<br>397-573 (177) | none | N/A |
| PML | PhaSePro | P29590-12 | 1-30 (30)<br>434-563 (130) | 49-104 (56)<br>119-167 (49) | 1bor.pdb<br>6imq.pdb |
| PSD-95 | PhaSePro | P78352-3 | 35-54 (20) | 55-243 (189)<br>302-399 (98) | 6spv.pdb<br>3i4w.pdb |
| AGO2 | PhaSePro | Q9UKV8 | none | 22-859 (838) | 4f3t.pdb |
| Matrin-3 | PhaSePro | P43243 | 38-111 (74)<br>125-232 (108)<br>245-282 (38)<br>331-400 (70)<br>588-790 (203) | none | N/A |
| U2AF65 | PhaSePro | P26368 | 1-89 (89)<br>113-145 (33) | 90-112 (23)<br>148-336 (189)<br>375-475 (101) | 1jmt.pdb<br>2g4b.pdb<br>4fxw.pdb |
| MORC3 | PhaSePro | Q14149 | 455-660 (206)<br>755-776 (22)<br>857-885 (29) | 9-454 (446) | 6o1e.pdb |
| YTHDF2 | PhaSePro | Q9Y5A9 | 1-48 (48)<br>217-394 (178) | 398-548 (151) | 4wqn.pdb |
| YTHDF1 | PhaSePro | Q9BYJ9 | 1-53 (53)<br>70-89 (20)<br>132-188 (57)<br>216-363 (148) | 364-558 (195) | 4rci.pdb |
| YTHDF3 | PhaSePro | Q7Z739 | 1-54 (54)<br>81-113 (33)<br>136-378 (243)<br>517-538 (22) | none | N/A |
| CPEB3 | PhaSePro | Q8NE35 | 1-372 (372)<br>407-433 (27) | 440-540 (101) | 2rug.pdb |
| 53BP1 | PhaSePro | Q12888 | 1-1247 (1247)<br>1260-1481 (222)<br>1621-1713 (93) | 1484-1603 (120)<br>1714-1972 (259) | 2g3r.pdb<br>1kzy.pdb |
| Amyloid-beta | PhaSePro | P05067 | 194-286 (93)<br>349-370 (22)<br>625-671 (47) | 28-123 (96)<br>287-342 (56)<br>371-566 (196)<br>672-699 (28) | 1mwp.pdb<br>1app.pdb<br>3nyl.pdb<br>1amb.pdb |
| UBQLN2 | DisProt | Q9UHD9 | 450-624 (175) | none | N/A |
| PAB1 | DisProt | P04147 | 419-503 (85) | none | N/A |

| | | | | | |
|---|---|---|---|---|---|
| mid1 | DisProt | P78953 | 1-452 (452) | none | N/A |
| RXRG | DisProt | P48443 | 2-127 (126) | none | N/A |
| EMB506 | DisProt | Q9SQK3 | 40-112 (73) | none | N/A |
| AKRP | DisProt | Q05753 | 37-231 (195) | none | N/A |

[a] List of phase separating proteins obtained from Vernon *et al* (20), the PhaSePro database (21), and IDPs annotated "liquid-liquid phase separation" (IDPO:00041) in the DisProt database (22). Duplicate entries were removed from the combined list. The proteins TAF15, FUS, EWS, DDX3X, DDX4, TIA1, Elastin, RBM14, ROA1, and ROA2, listed in Vernon *et al*, were removed from the PhaSePro human protein set. ROA1 and ROA2 were identified as HNRNPA1 and HNRNPA2B1, respectively, in PhaSePro. IDPs annotated "liquid-liquid phase separation" in DisProt and originating from FUS, laf1, ROA1, ROA2, and DDX4, listed in Vernon *et al*, and DAXX, p62, TDP43, Galectin-3, NPM1, and NCK1, listed in PhaSePro, were also removed from the combined set.

[b] ID regions ($N \geq 20$) were identified from sequence using the GeneSilico MetaDisorder Service that generates a consensus prediction based on 13 primary methods (23). When this service was not available, the IUPred2 long predictor (24) was used instead. PhaSePro already annotates proteins in its database for the presence of predicted IDRs, by using IUPred2, which we kept for our use here. In this list, the ID regions exclude residues that were verified as folded (see column 5; i.e., a position could not be classified as both ID and folded). IDRs obtained from DisProt were assumed to be fully ID, because DisProt is manually curated for verified cases of ID.

[c] Residue positions with resolved atomic coordinates in a PDB structure (x-ray or NMR) were used to verify regions ($N \geq 20$) that fold. Unresolved residue positions in a PDB structure were not classified as folded. Folded regions from different structures that overlapped were merged.

[e] The silk (spidroin-1) sequence at positions 77-96 is repeated at positions 477-496, 677-696, 877-896, 1277-1296, 1477-1496, 1677-1696, 1877-1896, 2077-2096, 2277-2296, 2477-2496 and 2677-2696. Only one copy of this sequence was kept in the testing set.

[f] The silk (spidroin-1) sequence at positions 122-259 is repeated at positions 322-459, 522-659, 722-859, 922-1059, 1122-1259, 1322-1459, 1522-1659, 1722-1859, 1922-2059, 2122-2259, 2322-2459, 2522-2659. Only one copy of this sequence was kept in the folded set.

**Table S3. Summary of Mann-Whitney U tests that compare mean $v_{model}$ (top) and mean β-turn propensity (bottom) in the null, testing, and folded sets.**

| $v_{model}$: | U test w/ null [a] | U test w/ folded [a] |
|---|---|---|
| null set | - | 2.5e-08 |
| testing set | 2.4e-04 | 7.9e-03 |
| folded set | 2.5e-08 | - |

| β-turn propensity: | U test w/ null [a] | U test w/ folded [a] |
|---|---|---|
| null set | - | 4.5e-08 |
| testing set | 1.0e-06 | <1.1e-16 |
| folded set | 4.5e-08 | - |

[a] one-tail p-value

**Table S4. Normalized frequency for β-turn.**

| Amino Acid | Scale value [a] |
|---|---|
| Alanine | 0.770 |
| Arginine | 0.880 |
| Asparagine | 1.280 |
| Aspartic Acid | 1.410 |
| Cysteine | 0.810 |
| Glutamine | 0.980 |
| Glutamic Acid | 0.990 |
| Glycine | 1.640 |
| Histidine | 0.680 |
| Isoleucine | 0.510 |
| Leucine | 0.580 |
| Lysine | 0.960 |
| Methionine | 0.410 |
| Phenylalanine | 0.590 |
| Proline | 1.910 |
| Serine | 1.320 |
| Threonine | 1.040 |
| Tryptophan | 0.760 |
| Tyrosine | 1.050 |
| Valine | 0.470 |

[a] From Levitt (25).

**Table S5. Structural properties of turn and non-turn ensembles.**

| | **Non-Turn Ensemble** [a] | **β-Turn Ensemble** [a] |
|---|---|---|
| Total ASA (Å$^2$) | 738.3 ± 0.9 | 670.3 ± 0.5 |
| Hydrophobic ASA (Å$^2$) | 536.4 ± 0.8 | 488.9 ± 0.5 |
| CHASA (Å$^2$) | 353.6 ± 0.6 | 327.3 ± 0.6 |
| Hydrophobic ASA lost assuming backbone hydration (Hydrophobic ASA – CHASA, Å$^2$) | 182.8 ± 1.0 | 161.6 ± 0.7 |
| Number of Backbone Hydration Waters (CHASA maximum is 55) | 44.4 ± 0.1 | 37.1 ± 0.1 |

[a] Uncertainties were calculated as the standard error of the mean.

**Table S6. List of proteins that exhibit phase separation behavior *in cellulo* that were found by *in vitro* characterization not to phase separate as purified proteins.**

| Name [a] | UniProt accession number |
|---|---|
| RBM3 | P98179 |
| ERF3 | P05453 |
| CPEB2 | Q7Z5Q1 |
| WASL | O08816 |
| FMR1 | Q06787 |
| LSM4 | P40070 |
| SynGap | J3QQ18 |
| SOS1 | Q07889 |
| PUB1 | P32588 |
| LAT | O43561-2 |
| Nephrin | O60500 |
| dcp2 | O13828 |
| pdc1 | O13892 |
| Disks4 | P78352 |
| GRB2 | P62993 |
| NCK1 | P16333 |
| edc3 | O94752 |
| npm1 | P07222 |

[a] List obtained from (20).

**Table S7. Summary of pair-wise Mann-Whitney U tests comparing the relative population (given by set percentage) of predicted PS region lengths, for lengths ranging from 1 to 150 residues.** The sequence sets that were compared are those described in Figure 5 in the main text. Here, we compare each set known to be enriched for LLPS (rows) to those sets that are not enriched for that property (columns). Reported values are one-tail p-values, where p-value < 0.05 indicates a statistically significant difference in the two distributions.

| | human proteome [a] | DisProt [b] | SCOPe [c] |
|---|---|---|---|
| in vitro LLPS sufficient | < 1.1e-16 | < 1.1e-16 | < 1.1e-16 |
| in vitro LLPS insufficient | 4.2e-12 | 7.2e-13 | < 1.1e-16 |
| DisProt LLPS annotated | < 1.1e-16 | < 1.1e-16 | < 1.1e-16 |
| PhaSePro | < 1.1e-16 | < 1.1e-16 | < 1.1e-16 |

[a] UniProt reference proteome UP000005640
[b] DisProt database minus LLPS annotated entries
[c] SCOPe database version 2.07

**Supporting Figures**

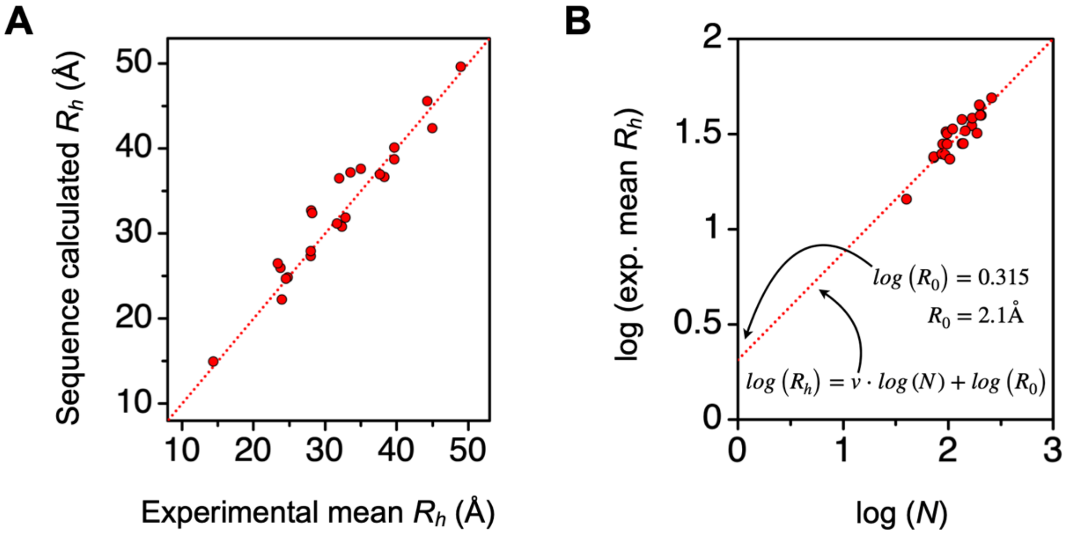


**Figure S1. Experimental mean $R_h$ compared to sequence calculated mean $R_h$. A)** The identity of the IDPs, sequences, and their experimental values are provided in Table S1. Sequence calculated mean $R_h$ was determined using equation [3], given in Experimental Procedures. The stippled line is the identity line. **B)** The y-axis intercept from the trend line (stippled line in figure) of a log-log plot of mean $R_h$ and $N$ (protein length) yields the pre-factor, $R_o$, in the power law scaling equation $R_h = R_o \cdot N^{\nu}$.

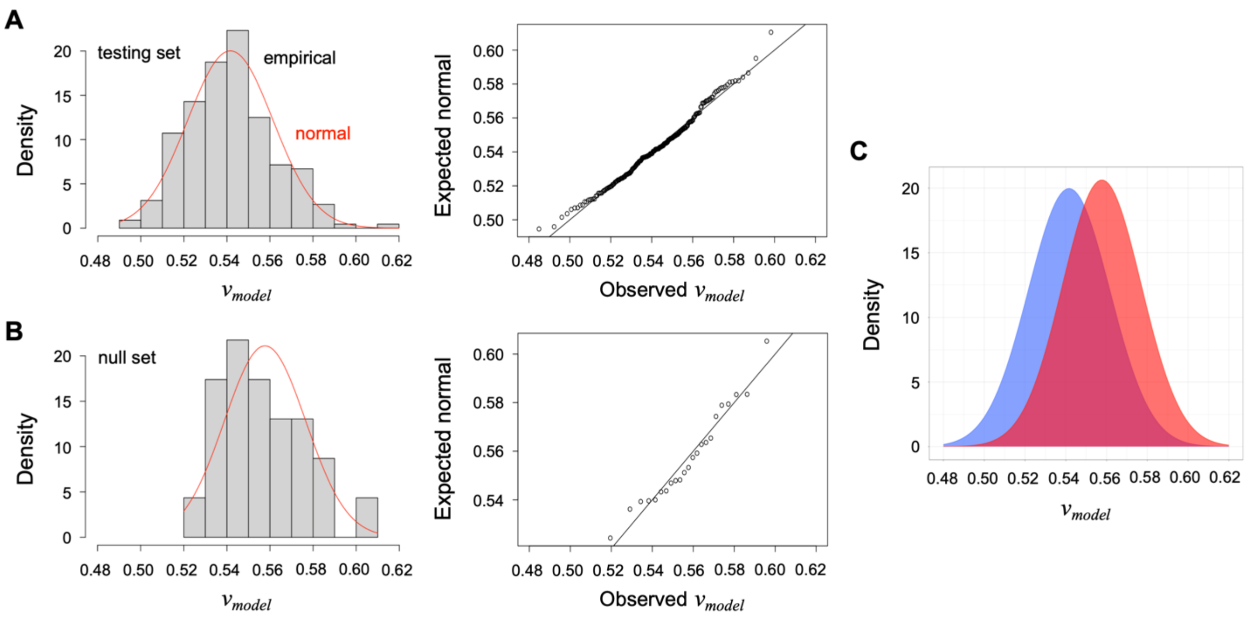


**Figure S2. Distribution of $v_{model}$ values in the A) testing and B) null sets.** The left-most figure in panels A and B compares the histogram distribution of $v_{model}$ values in the testing and null sets, respectively, to the probability density function of the normal distribution, $f(x) = \frac{1}{\sigma\sqrt{2\pi}} e^{-\frac{1}{2}\left(\frac{x-\mu}{\sigma}\right)^2}$, shown by the red line, where μ and σ are the distribution mean and standard deviation. The right-most figure is a Q-Q (quantile-quantile) plot that compares two probability distributions by plotting the quantiles against each other; in this case the observed empirical against the normal. When the trend in this plot follows the identity line (black line), this provides evidence that the compared distributions are similar. Because both the testing and null sets exhibit this behavior when compared to normal distributions, both data sets can be considered as similar to normal. Panel C overlays the distribution of $v_{model}$ values in the testing (blue) and null (red) sets when calculated as normal using the probability density function and their observed distribution mean and standard deviation.

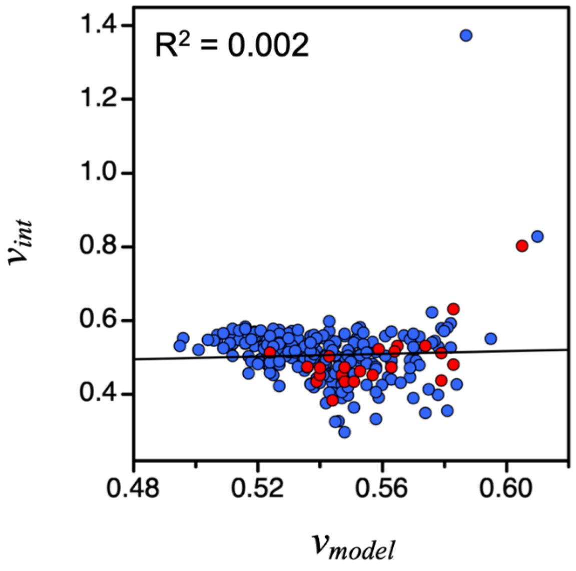


**Figure S3. Comparing sequence calculated $v_{model}$ and $v_{int}$.** Blue circles show the calculated values of $v_{model}$ and $v_{int}$ for the IDR sequences in the testing set (**Table S2**). Red circles show values for the IDP sequences in the null set (**Table S1**). The correlation, $R^2$, was calculated for the combined set of sequences, testing and null.

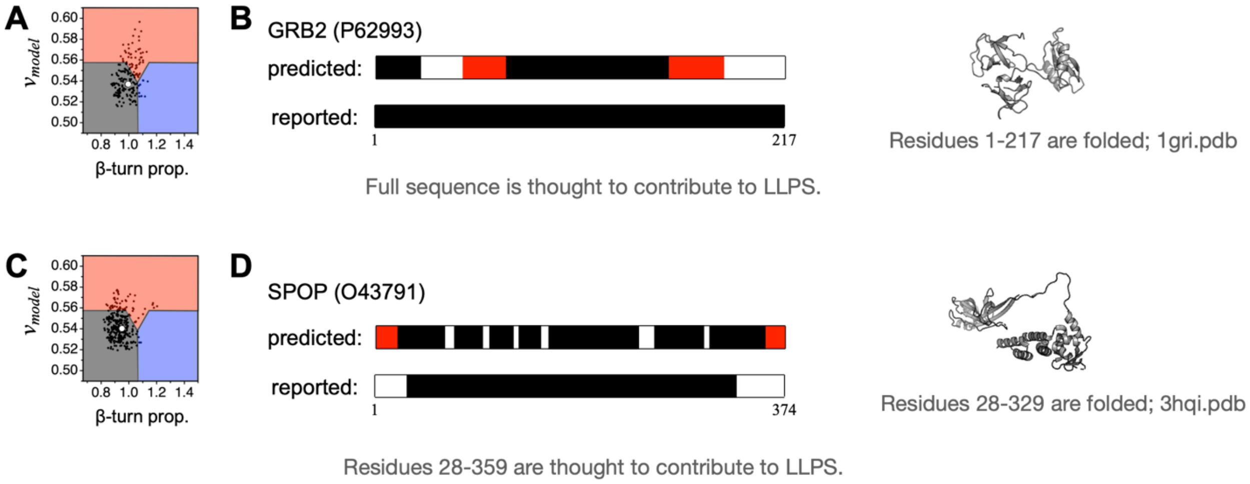


**Figure S4. Predicting protein regions that drive LLPS. A-D)** Using a sliding window algorithm applied to the primary sequence (described in Figure 4 legend of main text), contiguous regions ($N$≥20) that were 90% of only one label P, D, or F were colored blue, red, or black, respectively, to represent predicted PS, ID, or folded regions. The proteins in this figure are identified by name and UniProt accession number. The ParSe algorithm predicts there are no PS (i.e., phase separating ID) regions in these two proteins.

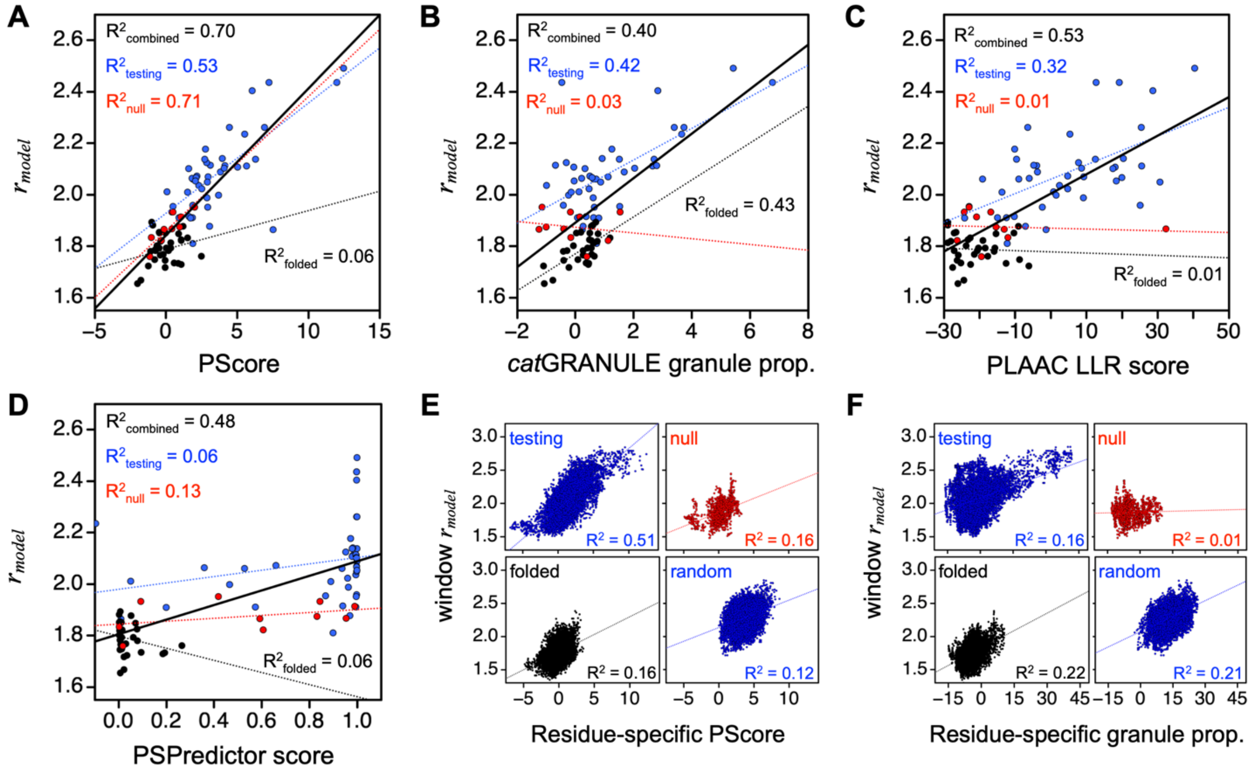


**Figure S5. Pair-wise correlations of predictor results.** Comparison of $r_{model}$ to **A)** PScore, **B)** granule propensity, **C)** LLR, and **D)** PSPredictor in the testing, null, folded, and combined sequence sets. Residue level comparison of window $r_{model}$, where the window value was assigned to the central residue position, to **E)** PScore and **F)** granule propensity for sequences in the testing (top left), null (top right), folded (bottom left), and randomized (bottom right) sets.
